# Convergent evolution of skipper wings assessed through anti-predator defences, flight proxies and geographical distribution

**DOI:** 10.1101/2025.06.23.661062

**Authors:** Daniel Linke, Vincent Debat, Pável Matos-Maraví

## Abstract

1. Phenotypic evolution is shaped by phylogenetic, ecological and biomechanical constraints. Butterfly wings offer an ideal system to explore these constraints as their evolution is strongly influenced by natural and sexual selection affecting colour patterns and wing shape in relation to visual signalling and flight performance.

2. We focus on Eudamina skippers, butterflies with large-scale phenotypic convergences involving wing shape and colour pattern. Using a comprehensive morphometric dataset encompassing 176 species, we assess their morphological evolution in a phylogenetic framework.

3. We show that hindwing tails and dorsal iridescence, proposed to signal high escape ability, have repeatedly evolved in tropical species. Hindwing tails are associated with intermediate body size, low wing loading and a forewing shape that facilitates manoeuvrability; iridescence is more common in large species with significantly high wing loading and aspect ratio, indicating high flight speed and manoeuvrability.

4. The evolution of all tested morphometric traits is best described by Ornstein-Uhlenbeck models, indicating stabilising selection around fitness optima in Eudamina. Models testing for random drift or early burst dynamics were rejected in all cases.

5. We reveal a clear co-evolution between the fore- and hindwing shapes, despite both wings having different functional constraints. Hindwing tails have a significant recurrent effect on forewing shape in lineages evolving them independently, suggesting the existence of evolutionary trade-offs between flight performance and antipredator defences.

6. Our study shows that the convergent morphologies across Eudamina have likely arisen not by neutral evolution but by ecological selection influenced by biomechanical constraints.

## Introduction

Phenotypic diversity is partly explained by the remarkable capacity of organisms to adapt to their environments, a process central to Darwinian evolution (Darwin, 1859). Phenotype evolution is constrained by the physical environment, biotic interactions, developmental mechanisms, and evolutionary history (Tendler et al., 2015; Galis et al., 2018; Borics et al., 2023). Despite the recognition of such constraints, our understanding of their influence and which potential trade-offs they impose on evolutionary pathways has remained largely qualitative for most animal groups (but see: Kang et al., 2017; Hamilton et al., 2022).

Butterflies provide an ideal system for investigating how wing phenotypes have evolved under ecological (Kang et al., 2017) and biomechanical constraints (Rossato et al., 2018; Le Roy et al., 2019; Hamilton et al., 2022). Butterfly wings are essential to survival and are constrained by different selective pressures related to flight performance, mate recognition, antipredation, and thermoregulation. Forewings are critical for powered flight, while hindwings—which are not strictly necessary for flight—play a significant role in evasive manoeuvres (Jantzen & Eisner, 2008). Fore- and hindwings thus experience different selective pressures that may translate to relative evolutionary independence. In line with this hypothesis, Owens et al. (2020) reported higher evolutionary rates in hindwing shape and size in the *Papilio* genus, interpreted as a stronger biomechanical constraint on the forewing. In contrast, Hamilton et al. (2022) reported a coevolution between the two wings, long hindwing tails being associated with simpler forewing shape in silk moths (Saturniidae: Arsenurinae). Developmental and biomechanical constraints might thus jointly affect the two wing pairs and rather promote their correlated evolution (Dudley, 2002). The degree to which fore- and hindwings of Lepidoptera co-evolve thus remains unclear and should be specifically quantified.

To better understand the factors shaping the evolution of butterfly wing shape and size, we focused on the Eudamina skippers (Lepidoptera: Hesperiidae: Eudaminae), a subtribe distributed across North and South America. Eudamina butterflies exhibit striking phenotypic variation, including multiple convergences of hindwing tails, dorsal iridescence and creamy bands on forewings (Li et al., 2019; Zhang et al., 2019, 2023). These convergent phenotypic traits have been hypothesized to signal unprofitability due to evasiveness (Janzen et al., 2009; Pinheiro & Freitas, 2014; Linke et al., 2022, 2024). Convergence among hard-to-catch prey is indeed expected if predators associate their visual appearance with difficulty of capture and avoid attacking them, a phenomenon referred to as evasive mimicry (van Someren & Jackson, 1959; Holling, 1965), which has received increased experimental attention recently (e.g., Páez et al., 2021; Linke et al., 2022; Loeffler-Henry & Sherratt, 2024). Dorsal iridescence has been suggested to signal evasiveness in skippers, due to its common expression in highly agile species (Janzen et al., 2009), although in other larger butterflies iridescence may work as a flash colouration, impeding predator attacks while the animal moves (Silvasti et al., 2024). Field observations with wild birds have also suggested that creamy bands on wings of highly evasive butterflies might signal flight prowess (Pinheiro & Freitas, 2014). Hindwing tails, on the other hand, might not only signal difficulty of capture (Linke et al., 2022), but also contribute to it in two ways (1) they might participate in aerodynamic performance, either in force production or in trajectory stabilization (Park et al., 2010) or (2) they might deflect predators attacks away from vital body parts. In *Iphiclides podalirius* (Papilionidae), captive birds indeed mostly attacked the tails of dummy butterflies in experimental trials (Chotard et al., 2022).

If evasive mimicry is indeed driving the convergent evolution of these different phenotypic traits, they should be tightly associated with the high escape capabilities they are presumed to signal, particularly those linked to flight performance. While flight performances are experimentally very challenging to measure, involving 3D videography of natural escape flights in natural conditions for many species, they can be assessed indirectly using morphometric proxies, that can be obtained for many specimens and species (for a review, see Le Roy et al., 2019). Wing loading (ratio of body weight to wing area) correlates with flight speed (Betts & Wootton, 1988), aspect ratio (elongation of the forewing) correlates with gliding capabilities (Cespedes et al., 2015) and second moment of the area (area distribution along the proximo-distal forewing axis) correlates with flapping efficiency (DeVries et al., 2010). While these proxies oversimplify the link between morphology and flight performance, they provide a valuable framework for phylogenetic comparative studies testing evolutionary relationships between ecological conditions and wing morphology.

We hypothesize that converging Eudamina wing phenotypes are associated with high flight capabilities and evolve under selection imposed by predation in tropical environments. In addition, a co-evolution of fore- and hindwing shapes is expected because of a joint selection related to aerodynamic performance. Through a combined morphometric and phylogenetic approach, we aim to test the following hypotheses:

1. If convergent phenotypes (hindwing tails, dorsal iridescence and creamy forewing bands) are indeed adaptive, we predict that their evolution will deviate from a Brownian-Motion model (supporting neutrality) and instead best fits an adaptive model (i.e., Early-Burst or Ornstein-Uhlenbeck).
2. If their evolution is not neutral, we expect that these convergent phenotypes are associated with elevated flight performances, as assessed through morphological flight proxies.
3. If hindwing tails have an impact on flight, we predict a coevolution with forewing shape, indicative of joint aerodynamic constraints on the two wing pairs.

## Material and Methods

### Butterfly images

We photographed the dorsal and ventral views of Eudamina butterflies archived at the Museum für Naturkunde in Berlin (19 specimens), the Natural History Museum in London (314 specimens), the Museo de Historia Natural in Lima (92 specimens) and the Biology Centre CAS in České Budějovice (4 specimens). We used a photo-box made of white plastic sheets (H = 50 cm, W = 30 cm, L = 30 cm) with LED lightning allowing for high colour accuracy (CRI > 95%). All photos were standardized using a grey background reference (18%, Color Balance Cards GC-1, JJC, Guangdong, China), which was complemented with a colour checker (colorchecker classic mini, calibrite, Wilmington, USA) and a metric scale. We used a Canon EOS 250D camera (Canon, Tokyo, Japan) with an EFS 18-55 mm lens and the photo parameters ISO = 100, F = 18, Tv = 1/20s (the shutter speed was increased to 1/40s in Lima, due to overexposed photos). To complement our photographic database, we retrieved photos including a metric scale from Butterflies of America (www.butterfliesofamerica.com, 39 specimens), published sources (Zhang et al., 2023; Bertrand et al., 2014; 9 specimens), and the Entomological collection in La Havana, Cuba (photos provided by Rayner Núñez of *Chioides marmorosa*, 3 specimens). Altogether, we utilised photographic data of 480 specimens from 176 out of the 179 valid Eudamina species (October 2023).

### Morphometrics measurements

Because of their overlap in mounted specimens, forewings and hindwings were measured using the dorsal and ventral sides, respectively. For area measurements, we used the histogram tool of GIMP v 2.10.24 (GNU Image Manipulation Program, Verden, Germany).

We used standard and comprehensive morphometric measurements in butterflies (Figure 1): **forewing length** (FWL, distance from the forewing’s base to the apical tip), **forewing width** (FWW, the maximum extend perpendicular to FWL), **thorax length** (TL, distance between the head and the abdomen), **thorax width** (TW, width of the thorax in the centre), **forewing area** (FWA), **hindwing area** (HWA), **hind wing length** (HWL, distance from the hindwing’s base to the end of vein Cu2), and **tail length** (distance from the hindwing’s base to the end of vein 2A).

**Figure 1:**
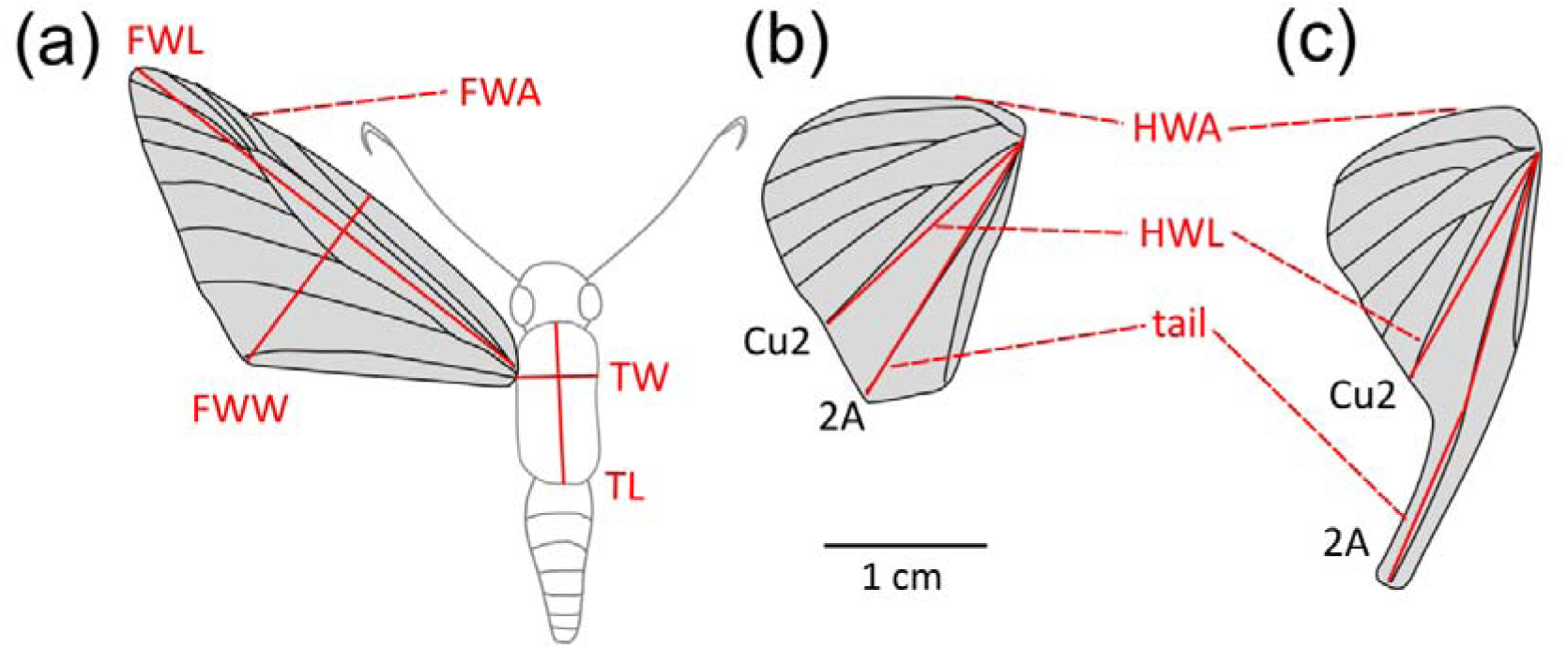
Butterfly morphometric measurements. (a) forewing and (b) hindwing of the tailless *Telegonus apastus*, and (c) hindwing of the tailed *Chioides catillus*. FWL = forewing length, FWW = forewing width, FWA = forewing area (in grey), TW = thorax width, TL = thorax length, HWA = hindwing area (in grey), HWL = hindwing length, tail = tail length. Hindwing veins used for measurements are highlighted (Cu2 and 2A).

We measured three morphological parameters associated with flight performance (Le Roy et al., 2019): **wing loading**, defined as the ratio of body mass to forewing area, **aspect ratio**, defined as the ratio of forewing length on forewing width, and **second moment of area**, defined as the distribution of area along a proximo-distal forewing axis. Wing loading and aspect ratio were derived from our morphometric measurements, following (Henriques et al., 2022; Kleckova et al., 2024). First, thoracic volume was estimated as 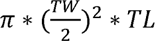, which was then used to calculate wing loading as 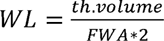. **Aspect ratio** was calculated as the ratio of FWL to FWW. The second moment of area was obtained using wingImageProcessor v.1.1 (www.biomech.web.unc.edu/wing-image-analysis) in MATLAB 24.1.0.2628055 (R2024a) Update 4 (MathWorks, Natick, USA). To obtain a relative measure of hindwing tail length compared to wing size, we computed **tail ratio**, as the proportion of tail length over hindwing length.

To precisely quantify the variation of wing shape, we used a geometric morphometric approach. Using tpsdig v. 2.32 (Rohlf, 2005) we defined 50 semi-landmarks on the outline of the forewing in dorsal view and of the hindwing in ventral view. The semi-landmark configurations were superimposed using *geomorph* v. 4.0.6 (Adams & Otárola-Castillo, 2013), minimizing the Procrustes distance during the sliding (Mitteroecker & Gunz, 2009). The coordinates after superimposition were used as shape variables in the subsequent multivariate analyses. The centroid size was also used as an estimator of wing size.

To create a comprehensive body size metric, we performed a principal component analysis (PCA) on highly correlated body size predictors, including HWA, HWL, FWL, FWW, FWA, TL, TW, thoracic volume and centroid size of the forewing shape, and used the first principal component (PC1) as our universal body size metric (explaining 90% of variability).

### Convergent traits

The different convergent traits (blue-green dorsal iridescence, creamy forewing bands and hindwing tails; Figure 2) were coded as binary traits (presence/absence). We defined a binary character state (presence/absence) for the tail, using a tail ratio cutoff of 1.3.

**Figure 2:**
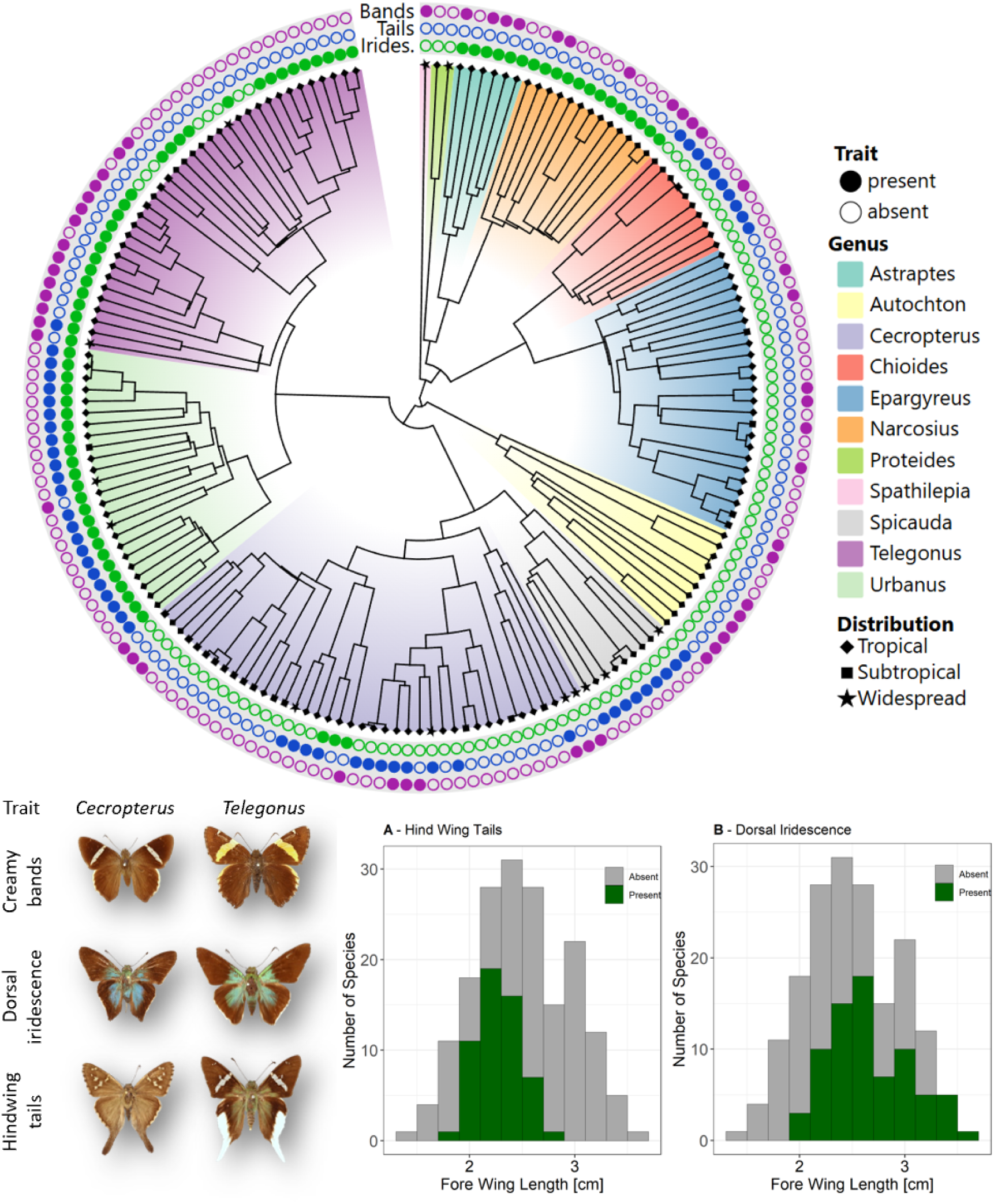
**Upper part**: Distribution of potential anti-predator defences along the phylogeny. **Lower left**: Representatives of species expressing creamy bands (*Cecropterus longipennis*, *Telegonus cellus*), dorsal iridescence (*Cecropterus egregius*, *Telegonus alardus*) and hindwing tails (*Cecropterus dorantes*, *Telegonus chalco*). **Lower right**: size dependant expression of anti-predator defences, **A** hindwing tails limited to intermediate sized species and **B** dorsal iridescence most common in larger ones.

### Distribution

Geographical distribution (tropics, subtropics, widespread) was scored for each species to evaluate how broad environmental variation affected butterfly wing evolution. We classified the geographical distribution of each species as centred in Neotropical biomes (Morrone, 2014), centred out of the Neotropics (i.e., subtropical), and widespread across tropical and subtropical North and South America. We determined the distributions using available distribution information from Butterflies of America (www.butterfliesofamerica.com), approximate range maps from the “Lepidoptera and some other life forms” database (www.nic.funet.fi/pub/sci/bio/life/insecta/lepidoptera/) and the geographical metadata gathered from collections.

### Phylogenetic inference

To infer phylogenetic relationships, we retrieved all available mitochondrial COI sequences of Eudamina from unpublished genome sequences (Matos-Maraví et al., *in prep.*), published data (Li et al., 2019; Ribeiro et al., 2021) and public databases (BOLD, www.boldsystems.org/ and NCBI, www.ncbi.nlm.nih.gov/) (Appendix S1: Table S1), obtaining genetic information for 139 species. We used BEAST v2.7.6 (Bouckaert et al., 2014) to time calibrate the phylogeny of Eudamina using a relaxed clock model and secondary calibration points as normal prior distributions based on Kawahara et al. (2023) on Eudamina’s stem (95% HPD: 19.59 - 23.95) and crown nodes (95% HPD: 15.19 - 19.15). We added all remaining missing species of Eudamina in the time-calibrated phylogeny by jointly modelling their evolution (without sequence data) within their respective genera, and further constrained inter-generic relationships based on published butterfly phylogenomic trees (Li et al., 2019; Kawahara et al., 2023). We ran four independent analyses, with sampling every 5,000 generations for 50 million generations. We combined all posterior trees and checked that the ESS values were higher than 200, before summarizing a maximum clade credibility (MCC) tree (Appendix S2: Figure S1).

All statistics use species level averages. Statistics were computed using R v. 4.3.0 (R Core Team, 2022), graphics were created using *ggplot2* (Wickham, 2016).

Ancestral state reconstructions (ASR) of convergent traits (hindwing tails, dorsal iridescence, creamy bands), body size and morphometric flight proxies (wing loading, second moment of area, aspect ratio) were performed using the *‘fastAnc’* function (*phytools* v. 1.5-1; Revell, 2012), and we visualised the trait evolution across the MCC tree by creating a continuous trait map using *‘contMap’* function of *phytools* v. 1.5-1 (Revell, 2012).

### Assessing phylogenetic signal and evolutionary models

We estimated Pageĺs lambda (λ) and Blomberǵs K for selected morphometric parameters (body size, wing loading, second moment of area, aspect ratio and tail ratio) using the ‘*phylosig*’ function of *phytools* v. 1.5-1 (Revell, 2012) and the MCC tree. The phylogenetic signal of the fore- and hindwing shapes (K_mult_) were calculated using the *‘physignal’* in the *geomorph* package v. 4.0 (Adams & Otárola-Castillo, 2013).

To test our first hypothesis, that Eudamina convergent traits are adaptive and have not evolved neutrally, we compared three evolutionary models, Brownian Motion (BM, ‘*mvBM*’), Ornstein-Uhlenbeck Model (OU, ‘*mvOU*’) and Early Burst (EB, ‘*mvEB*’) using the mvMORPH package v.1.2.1 (Clavel et al., 2015). BM assumes a constant neutral evolution of traits across linages, OU assumes the presence of an adaptive optimum, and EB assumes an initially high rate of trait evolution which slows down through time. For wing shapes, we performed a PCA and used the first 4 PCs for testing the three competing evolutionary models; they explained 92.5% variance for the forewing and 96.9% for the hindwing shapes. The model with the lowest AICc value (corrected Akaike information criterion) was chosen as the best fitting model. In addition to the MCC tree, we estimated phylogenetic signal and compared the fit of the three competing evolutionary models (BM, OU, and EB) across 100 randomly selected trees to assess the consistency of our results accounting phylogenetic uncertainty. Trees resulting in at least one non-converging model were removed from this comparison. To ensure equal variance and comparability among morphometric measurements, we scaled each parameter before fitting the evolutionary models (shape PCs were not scaled).

### Association between convergent traits and proxies of flight performance

To test whether convergent phenotypic traits (creamy bands, iridescence and tails) are linked to high flight abilities (assessed using morphometric proxies), we used phylogenetic Bayesian MCMC (Markov chain Monte Carlo) regression models due to their flexible hierarchical and phylogenetic modelling (4 chains, 4,000 iterations each, adapt delta set to 0.99 and max tree depth set to 10) using the R package *brms* v. 2.19.2 (Bürkner, 2017). To account for phylogenetic relatedness, a variance-co-variance matrix was calculated using the MCC tree in *ape* v. 5.7-1 (Paradis & Schliep, 2019). The variations of all continuous variables (responses and predictors) were scaled to assure equal variances. Our predictors in all models were: presence/absence of creamy bands and dorsal iridescence, tail ratio and geographical distribution (Tropical, Subtropical, Widespread). The phylogenetic variance-covariance matrix was treated as a random effect. As responses, we used our body size metric (PC1) and the three relevant flight proxies: wing loading, forewing second moment of area and aspect ratio. To mitigate multicollinearity among the morphometric flight proxies, we therefore ran four separate models, with the same set of predictors but a different response variable.

To visualise wing shape variation and phylogenetic relationships across Eudamina, we generated a phylomorphospace using the *phylomorphospace* function of *phytools* v. 2.3-0. (Revell, 2012). Relationships between wing shapes and convergent traits (hindwing tails, dorsal iridescence and creamy bands) were tested using Procrustes phylogenetic generalized least squares (PGLS) tests using the *‘procD.pgls’* function from the *geomorph* package (Adams & Otárola-Castillo, 2013). The MCC tree was used to account for shared evolutionary history among species, ensuring that relationships were assessed while accounting for phylogenetic non-independence. Separate models were run for forewing (FW) and hindwing (HW) shapes as responses. As predictors, we used body size (PC1), tail ratio, iridescence (presence/absence), creamy bands (presence/absence) and distribution (Tropical, Subtropical, Widespread). Tail ratio was not included in the model assessing hindwing shape, as this was redundant. Significance was determined by 1,000 permutations for each model.

To evaluate the covariation between fore- and hindwing shapes, we conducted a two-block partial least squares (PLS) analysis using the *‘two.b.pls’* function from the *geomorph* package (Adams & Otárola-Castillo, 2013). Phylogenetic PLS analysis was also run to assess the influence of phylogenetic relatedness on the co-variation, using the *‘phylo.integration’* function from *geomorph* (Adams & Otárola-Castillo, 2013). *Chioides vintra* was excluded from this analysis due to its unusually short genetic distance from its sister species, *Chioides catillus*, combined with an extremely large phenotypic divergence between both species, which disrupted the analysis. To better understand the impact of tails on flight performance, we specifically assessed the effect of tail length on the shape of the forewing with a multivariate regression of tail-ratio on forewing shape, using a custom-written R function (Appendix S3).

## Results

### Repeated evolution of tails and iridescence in the tropics

Hindwing tails were most common in widespread (53.8%, 7 of 13 species) and tropical species (32.9%, 46 of 140 species) while being least common in subtropical ones (8.7%, 2 of 23 species). Dorsal iridescence was most common in species with tropical distribution (49.3%, 69 of 140 species) while being considerably less common in widespread and subtropical species (23.1%, 3 of 23 species and 8.7%, 2 of 23 species, respectively). Lastly, creamy bands showed almost equal prevalence among regions (Subtropical: 26.1%, 6 of 23 species; Tropical: 30.7%, 43 of 140 species; Widespread: 30.8%, 4 of 13 species). Dorsal iridescence and hindwing tails were more prevalent at specific sizes: while iridescence was most common in the intermediate to large species, tails were most common in intermediate to low sized species (Figure 2A+B). Creamy bands were expressed equally across the size spectrum.

Ancestral state reconstruction suggested that hindwing tails evolved independently in four lineages (*Urbanus*, *Chioides*, *Telegonus*, and the common ancestor of *Spicauda* and *Cecropterus*), with multiple gains and losses within *Cecropterus*, particularly in subtropical species. Dorsal iridescence evolved independently in *Cecropterus* and in the common ancestor of *Narcosius* and *Astraptes* as well as of the clade consisting of *Urbanus* and *Telegonus*, in which it was lost in several times. Creamy bands were more evenly distributed across the phylogeny with no clear pattern. Plots of the ancestral state reconstruction of tail ratio, iridescence, bands, body size (PC1), wing loading, aspect ratio and second moment of area can be found in Appendix S4: Figure S3-S9.

### Phenotypic traits evolution follows OU models in Eudaminae

For body size (PC1), there was a strong phylogenetic signal (Pagel’s lambda = 0.866; Blomberg’s K = 0.529) and the Ornstein-Uhlenbeck model provided the best fit (AICc = 392.703), indicating a stabilising selection towards trait optima for body size. Wing loading, aspect ratio, second moment of area and tail ratio had phylogenetic signals of varying strength (λ = 0.635, 0.838, 0.001 and 0.909; K = 0.186, 0.359, 0.098 and 0.281, respectively) and OU was again consistently selected as best (AICc = 504.983, 427.987, 506.186 and 468.804; ΔAICc > 50 vs. BM or EB in each case), confirming stabilising selection on those traits. Shapes of fore- and hindwings were also best explained by the OU model (AICc = −3432.267 and −2302.417, respectively), with significant phylogenetic influence on shape evolution (Kmult = 0.202 and 0.316). Results averaged over 100 posterior trees (N ≈ 50–90) closely match the MCC-tree values, demonstrating that OU processes dominate trait evolution in Eudamina.

### Tails and iridescence are associated with different flight performance proxies

Testing for associations between morphological measurements used as proxies for flight performance and convergent traits using brms models revealed several key relationships. In the first model using size (PC1) as a response, tail ratio showed a slightly negative association with body size, indicating that species with hindwing tails tend to be smaller than those without (mean = −0.14, SD = 0.09, L-95% CI = −0.31, U-95% CI = 0.04, see Figure 1A). Widespread species tend to be significantly larger than subtropical ones (widespread compared to subtropical; mean = 0.44, SD = 0.22, L-95% CI = 0.01, U-95% CI = 0.88), further the model suggested that tropical species tend to be larger than subtropical ones, although with low probability considering the posterior intervals (mean = 0.25, SD = 0.14, L-95% CI = −0.02, U-95% CI = 0.52).

In the second model testing the effects on wing loading, the presence of iridescence was significantly associated with increased wing loading (mean = 0.94, SD = 0.24, L-95% CI = 0.46, U-95% CI = 1.41), suggesting a fast flight in species expressing dorsal iridescence. Additionally, a significant negative effect of tail ratio was present (mean = −0.19, SD = 0.09, L-95% CI = −0.38, U-95% CI = −0.01) indicating lower wing loading, i.e., lower flight speed, in tailed compared to tailless species. All other predictors showed no strong effect based on the estimated intervals.

The third model tested the effects on second moment of area (a proxy for flapping efficiency). While no comparison yielded a strong statistical effect, the results suggested a low second moment of area in tailed species (mean = −0.11, SD = 0.09, L-95% CI = −0.29, U-95% CI = 0.06), indicating a limited flapping flight power but high manoeuvrability (Betts & Wootton, 1988). A higher second moment of area in species with tropical distribution compared to subtropical ones is also detected (mean = 0.43, SD = 0.24, L-95% CI = −0.05, U-95% CI = 0.90).

The fourth model testing the effects on aspect ratio only revealed a strong positive association with the presence of iridescence (mean = 0.49, SD = 0.23, L-95% CI = 0.05, U-95% CI = 0.93). Iridescent species were therefore associated with high wing loading and high aspect ratio, a combination likely associated with fast sustained flight (Betts & Wootton, 1988). The complete model outputs and diagnostic plots can be found in Appendix S5. The Bayesian MCMC regression analyses exhibited strong convergence diagnostics, with potential scale reduction factors (Rhat) consistently equal to 1 and effective sample sizes exceeding 3,700 for all parameters, indicating that the Markov chains mixed well and that the posterior estimates are robust and reliable.

To visualise the variation in fore- and hindwing shape, we generated a phylomorphospace, combined with density curves (Figure 3). As expected, the hindwing shape of tailed and tailless species forms two separate clusters. For forewings, while a slight shift along PC2 is detected suggesting a different forewing shape depending on the presence of tails on the hindwing, the clearest pattern is the reduced forewing shape variation of tailed species, along PC1, possibly indicative of aerodynamic constraint.

**Figure 3:**
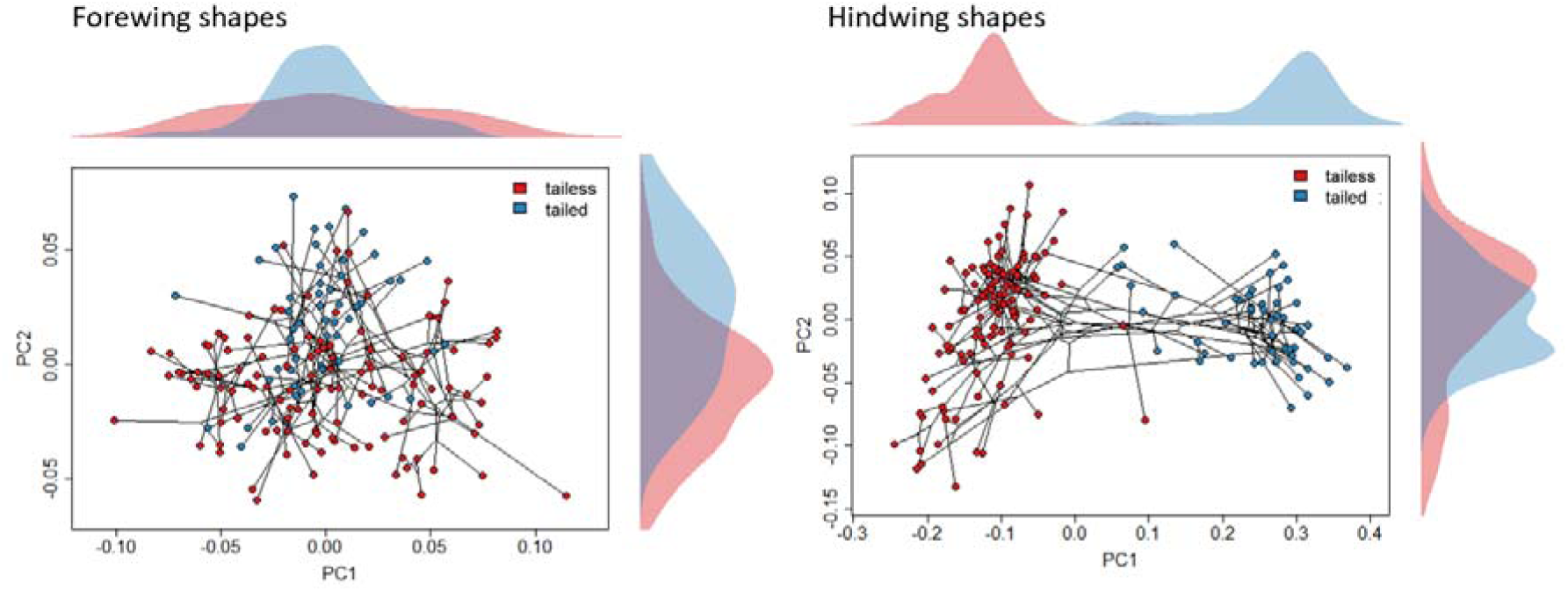
Phylo-morphospace for fore-(left) and hindwings (right), indicating the presence / absence of hindwing tails (present at tail ratios above 1.3)

When testing the effect of body size, convergent traits and geographical distribution on wing shape variation using a PGLS model, nearly all parameters had significant effects. For forewing shapes, the PGLS model showed significant effects of body size (PC1) (F = 19.68, p = 0.001), geographical distribution (F = 7.98, p = 0.003), creamy band presence (F = 12.34, p = 0.002), tail ratio (F = 6.86, p = 0.005), and dorsal iridescence presence (F = 14.22, p = 0.001). For hindwing shape, body size (F = 13.780, p = 0.001), creamy bands (F = 4.614, p = 0.028) and dorsal iridescence (F = 6.459, p = 0.025) had significant effects. Geographical distribution had a marginal yet not significant effect (F = 1.586, p = 0.095, Table 2). For all significant factors, the explained variance (R^2^) was relatively low between 0.023 and 0.083. While body size had the greatest effect on wing shapes, the presence of traits associated with evasiveness did also affect fore and hindwing shapes significantly.

**Table 1:** Evolutionary model fit statistics, AICc and in brackets loglikelihood and phylogenetic signals (Pagel’s lambda, Bloomberg’s K, Kmult) for morphological traits, including body size, wing loading, second moment of area, aspect ratio, tail ratio, and fore- and hindwing shapes. Results for the MCC tree as well as 100 random trees are included (AVG ± SD). Best-fit evolutionary models are marked in **bold**.

| Trait | Phylogeny | Pagel's lambda | Blomberg's K | Kmult | Brownian Motion | Ornstein-Uhlenbeck | Early Burst |
| --- | --- | --- | --- | --- | --- | --- | --- |
| **Body Size (PC1)** | MMC | 0.866<br>(-339.815) | 0.529 |  | 427.560<br>(-211.764) | **392.703**<br>**(-192.234)** | 429.668<br>(-211.764) |
| | 100 trees<br>(N = 77) | 0.847 ±<br>0.022 | 0.197 ±<br>0.103 | | 581.466 ±<br>148.303 | **462.021 ±**<br>**35.741** | 583.536 ±<br>148.303 |
|  |  | (-155.391 ± 2.360) |  |  | (-288.698 ± 74.151) | **(-227.941 ± 17.871)** | (-288.698 ± 74.151) |
| **Wing loading** | MMC | 0.635 (612.686) | 0.186 |  | 608.943 (-302.437) | **504.983 (-248.375)** | 611.013 (-302.437) |
|  | 100 trees (N = 77) | 0.847 ± 0.022 (-155.391 ± 2.360) | 0.197 ± 0.103 |  | 581.466 ± 148.303 (-288.698 ± 74.151) | **462.021 ± 35.741 (-227.941 ± 17.871)** | 583.536 ± 148.303 (-288.698 ± 74.151) |
| **Second moment of area** | MMC | 0.001 (502.704) | 0.098 |  | 722.056 (-358.993) | **506.186 (-248.976)** | 724.126 (-358.993) |
|  | 100 trees (N = 77) | 0.001 ± 0.000 (-249.233 ± 0.000) | 0.055 ± 0.023 |  | 783.904 ± 100.681 (-389.918 ± 50.341) | **502.429 ± 1.777 (-248.145 ± 0.888)** | 785.975 ± 100.681 (-389.918 ± 50.341) |
| **Aspect ratio** | MMC | 0.838 (182.528) | 0.359 |  | 492.921 (-244.426) | **427.987 (-209.876)** | 494.991 (-244.426) |
|  | 100 trees (N = 89) | 0.786 ± 0.039 (-160.172 ± 2.671) | 0.134 ± 0.077 |  | 652.336 ± 158.733 (-324.133 ± 79.366) | **484.388 ± 22.538 (-239.124 ± 11.269)** | 654.406 ± 158.733 (-324.133 ± 79.366) |
| **Tail ratio** | MMC | 0.909 (62.844) | 0.281 |  | 538.524 (-267.228) | **468.804 (-230.285)** | 540.594 (-267.228) |
|  | 100 trees (N = 85) | 0.915 ± 0.013 (-128.869 ± 4.839) | 0.179 ± 0.155 |  | 643.016 ± 215.068 (-319.473 ± 107.534) | **462.659 ± 49.933 (-228.260 ± 24.966)** | 645.086 ± 215.068 (-319.473 ± 107.534) |
| **Forewing shape** | MMC |  |  | 0.202 | -3069.145 (1548.877) | **-3432.267 (1745.336)** | -2900.639 (1465.668) |
|  | 100 trees (N = 53) |  |  | 0.058 ± 0.037 | -2496.748 ± 330.352 (1262.679 ± 165.176) | **-3476.795 ± 59.749 (1763.281 ± 29.875)** | -2473.538 ± 319.936 (1252.118 ± 159.968) |
| **Hindwing shape** | MMC |  |  | 0.316 | -1987.302 (1007.956) | **-2302.417 (1180.411)** | -1871.161 (950.930) |
|  | 100 trees (N = 66) |  |  | 0.143 ± 0.108 | -1621.344 ± 312.677 (824.977 ± 156.339) | **-2390.663 ± 76.320 (1220.215 ± 38.160)** | -1613.242 ± 307.987 (821.970 ± 153.993) |

**Table 2:** Results of PGLS models for shape variation: Effects of anti-predator defences, geographical distribution and body size.

| Response | Predictor | Df | Sum of squares | R-squared | F-value | Z-value | p-value |
| --- | --- | --- | --- | --- | --- | --- | --- |
| Forewing shape | Body size (PC1) | 1 | 0.013 | 0.083 | 19.678 | 3.730 | 0.001 |
|  | Distribution | 2 | 0.011 | 0.067 | 7.984 | 2.413 | 0.003 |
|  | Bands | 1 | 0.008 | 0.052 | 12.341 | 2.738 | 0.002 |
|  | Tail ratio | 1 | 0.005 | 0.029 | 6.684 | 2.568 | 0.005 |
|  | Iridescence | 1 | 0.009 | 0.060 | 14.217 | 4.159 | 0.001 |
| Hindwing shape | Body size (PC1) | 1 | 0.083 | 0.070 | 13.780 | 2.408 | 0.013 |
|  | Distribution | 2 | 0.019 | 0.016 | 1.586 | 1.222 | 0.095 |
|  | Bands | 1 | 0.028 | 0.023 | 4.614 | 1.899 | 0.028 |
|  | Iridescence | 1 | 0.039 | 0.033 | 6.459 | 2.085 | 0.025 |

The two-block partial least squares analysis (PLS) on fore- and hindwing shapes revealed a strong correlation between both wings with an r-PLS of 0.538 (p = 0.001, Z = 4.90). When accounting for phylogenetic relationships using a phylogenetic PLS model (excluding *Ch. vintra*), the correlation coefficient was even higher (r-PLS = 0.648, Z = 5.75, p = 0.001), suggesting a strong covariation between fore- and hindwing shapes (Figure 4, Appendix S6: Figure S10), and showing that the two wings do not evolve independently.

**Figure 4:**
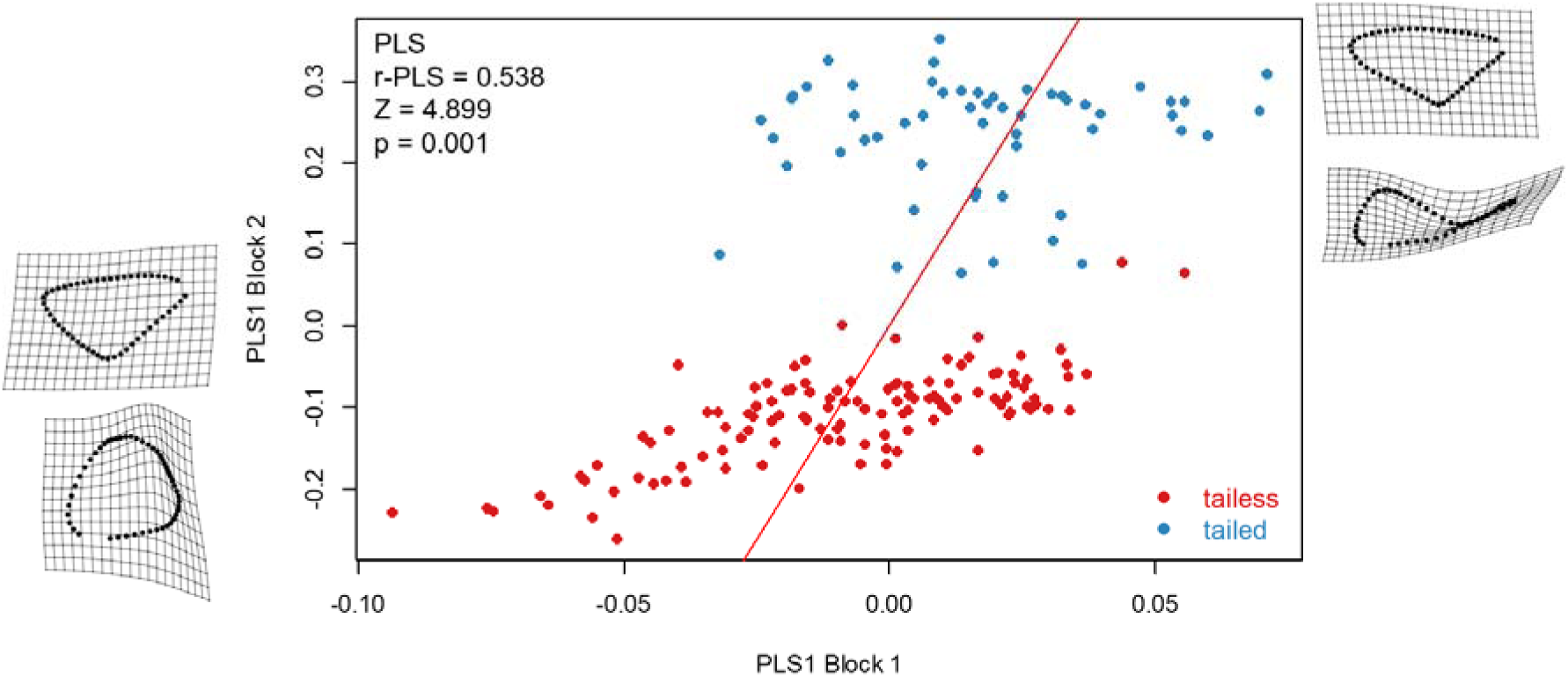
PLS analysis showing covariation between forewing and hindwing shapes. Wing shapes (forewings on x-axis, hindwings on y-axis) in the corner represent expected shapes in the extremes of their variation. Colour indicates the presence and absence of hindwing tails (presence = tail ratio > 1.3). Phylogenetic PLS plot can be found in Appendix S5: Figure S10.

Lastly, to assess the effect of hindwing tails on forewing shape, we applied a multivariate regression of forewing shape on tail ratio, allowing to reconstruct the expected forewing shape at extreme tail ratios (Figure 5). Reconstructed forewing shapes showed recurrent changes especially at the costal margin which was convex in species lacking hindwing tails and concave in species expressing tails. Additionally, the wing apex was narrower and the wing base broader in tailless compared to tailed species.

**Figure 5:**
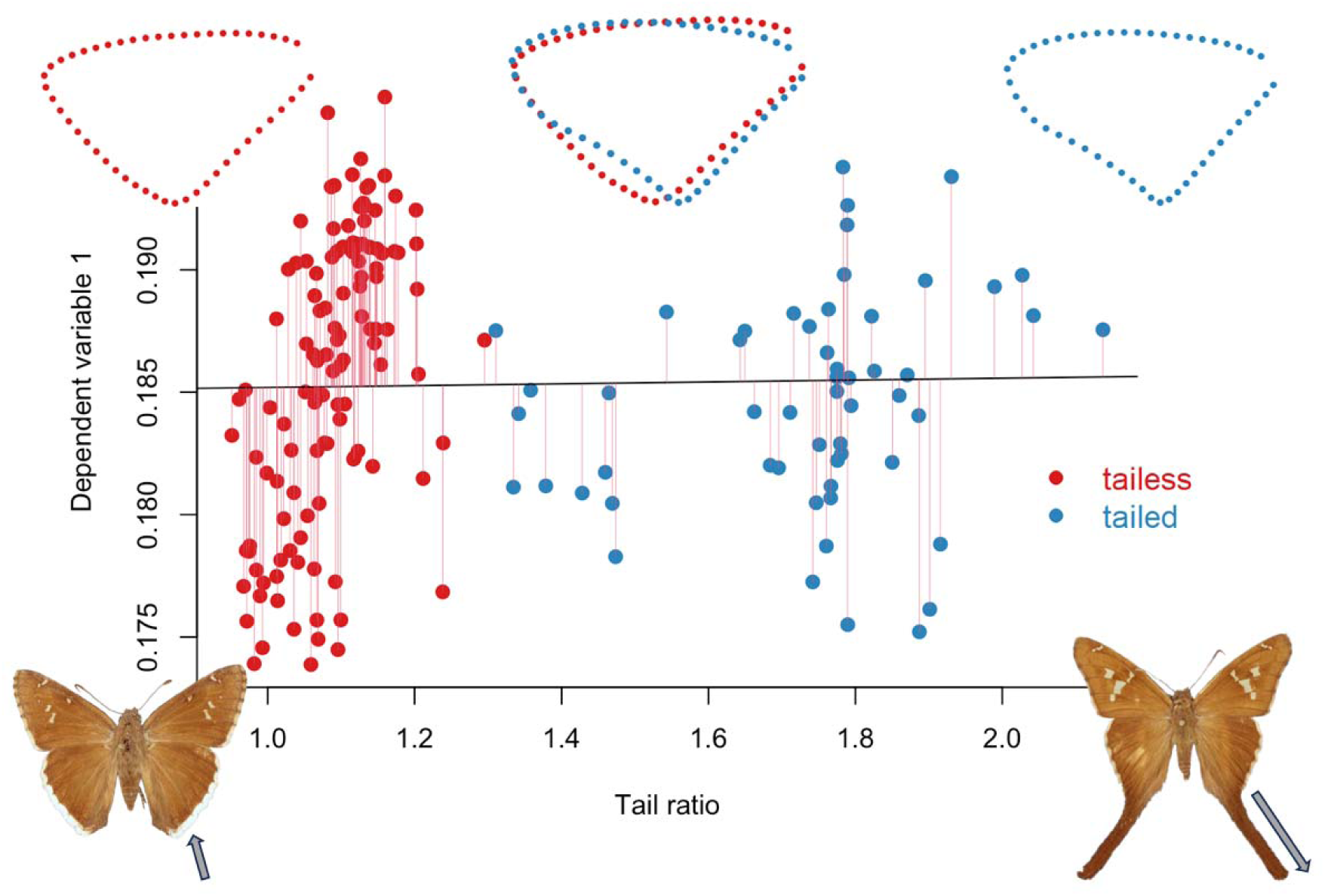
Visualising the multivariate regression of forewing shape on tail ratio, including regression curve and residuals. Wing shapes on top represent forewing shape at: blue = maximum observed tail ratio and red = minimum observed tail ratio (difference in shape amplified x2). Bottom left: *Cecropterus casica*, right: *Chioides albofasciatus*.

## Discussion

Our study shows evolutionary associations between convergent traits across skipper species (iridescence and the presence of hindwing tails), morphometric flight proxies and wing shapes indicative of high speed and manoeuvrability, suggesting that these traits might be associated with high escape abilities. If predators avoid attacking such hard-to-catch prey on the basis of these visual cues, these convergences among distantly related species might thus be driven by selection related to predation, a phenomenon dubbed evasive mimicry (van Someren & Jackson, 1959; Holling, 1965).

### Do wing tails signal, enhance or reduce escape performance?

The evolutionary drivers of wing tails in butterflies are remarkably poorly known considering their conspicuousness and their frequency, occurring in all butterfly families. While some evidence supports an effect of tails on predator attack deflection in Lycaenidae (Robbins, 1981; reviewed in Hendrick et al., 2022) and Papilionidae (Chotard et al., 2022), alternative possibilities like their effect of flight remain poorly investigated (but see: Park et al., 2010). Experimental work in captivity has shown that birds can use tails as a visual cue for difficulty of capture when associated with (artificial) escape prowess (Linke et al., 2022), however it is unknown whether tails indeed indicate high escape ability in nature. It is also unclear how the presence of tails might directly affect flight speed and escape performance. A study of butterfly wing aerodynamics, using metal models in a wind-tunnel, has shown that tails may enhance flight performance by increasing gliding flight stability (Park et al., 2010). In contrast, experiments with live moths with modified tails, either removed or artificially elongated, failed to detect any effect on flight aerodynamics (Rubin et al., 2018). For Eudamina, we showed that hindwing tails are significantly associated with decreased wing loading, which is typically linked to flight speed (Betts & Wootton, 1988; Le Roy et al., 2019). However, estimated wing loadings from widespread tailed skippers species (*Cecropterus dorantes*, *Spicauda simplicius*) are still higher than the majority of other studied butterfly species (Linke et al., 2024). While tailed species have lower wing loading, and by extension potentially lower flight speeds relative to other skippers, this does not imply they are easy to catch. The significant effect of tail-ratio on forewing shape suggests that tails indeed impact flight. In their influential study on how of wing shape modulates flight performance in butterflies, Betts and Wootton (1988) predicted that “*forewings with extended, narrow tips would gain some of the benefits of a high aspect ratio while maintaining reasonably low moments of area and mass, and hence high agility*”. This description matches the forewing shape associated with long tails we obtain in the multivariate regression (Figure 5), and the reduction of second moment of area associated with tails. This suggests that tails are associated with forewing shapes favouring a high agility.

Two competing hypotheses can thus be proposed: (1) if tails are involved in attack deflection, their presence might hinder flight performance, leading to selection for specific forewing shapes that compensate for this. (2) Conversely, tails may enhance flapping flight by improving manoeuvrability, favouring forewing shapes that support this. These hypotheses can be tested through the comparative analysis of flight trajectories and kinematics in both tailed and untailed species from different butterfly families expressing them, e.g., Papilionidae, Lycaenidae and Hesperiidae, or by comparing flight trajectories before and after tail loss.

This coevolution of the shapes of the two pairs of wings contradicts previous findings in Papilionidae, where fore and hindwings were suggested to evolve independently (Owens et al., 2020). This joint evolution of the two wings, but also of wing shape and colour patterns, underlines the integrated effects of selective pressures imposed by predators on the wing phenotypes.

The expression of tails in small to intermediate sized species, a pattern also found in silk moths (Hamilton et al., 2022), suggests potential negative trade-offs between body size and hindwing tails, possibly due aerodynamics constraints. In particular, the lack of tails in large and heavy species presenting a high wing loading – and thus high speed – point at a possible trade-off between flight power (high wing loading and no tail) and agility (low wing loading, presence of tails).

### Iridescence advertising fast flight?

The presence of iridescence (42% of all sampled species) is positively correlated with increased wing loading and aspect ratio in phylogenetic tests. Wing loading is positively correlated with increased flight speed (Le Roy et al., 2019), while aspect ratio has largely been linked to gliding abilities in Nymphalidae (DeVries et al., 2010; Cespedes et al., 2015; Le Roy et al., 2019). However, gliding behaviour is not known for skippers. Alternatively, aspect ratio has been linked to acceleration capability in male *Parage aegeria*, Nymphalidae (Berwaerts et al., 2002) or increased flight endurance and dispersal capabilities in other Lepidoptera (Qing & Zhi-tao, 2001; Jyothi et al., 2021; Jahant-Miller et al., 2022) suggesting a broader association with flight performance. That both metrics correlate with the presence of dorsal iridescence supports the hypothesis raised by Janzen et al. (2009), that iridescence indeed signals escape ability – and in particular flight power – in skippers.

Iridescence is mostly dorsal on butterfly wings and so has often been discussed in the context of flash colouration in large species such as *Morpho*, where the alternate exposure of the bright and dull wing surfaces induces colour changes during wing beats, that confuse predators and enhance the difficulty of capture (Murali, 2018; Silvasti et al., 2024; Vieira-Silva et al., 2024). However, due to their small size and high wing beat frequency, it remains unclear whether flash colouration can work for skippers. If iridescence contributes to the difficulty of capturing skippers, the lack of iridescence in small Eudamina species suggests that a relatively large size in skippers may be required for iridescence to have any anti-predator effect. Alternatively, iridescence might signal unpalatability. It has been shown that large skipper species with high wing loading are unpalatable to naïve predators (Linke et al., 2024), but the study only included one large iridescent species (*Telegonus fulgerator*). It is therefore unknown whether potential chemical defences leading to unpalatability are also common in other iridescent species.

### Creamy bands evolve neutrally

Creamy bands on forewings (30% of all sampled species) are not associated with specific flight proxies, geographical distributions or specific lineages. The lack of significant interactions between the expression of bands and morphological flight proxies does not suggest any link with flight performance. This finding contradicts the hypothesis that creamy bands might signal evasiveness in Neotropical butterflies (Pinheiro & Freitas, 2014). While no sexual dimorphism is present within Eudamina, it should be noted that the expression of creamy bands is largely limited to females in the tribe Phocidini (Eudaminae), e.g., the genera *Bungalotis* and *Nascus*, but their selective mechanism is unclear.

### Macro-evolutionary patterns support adaptive evolution of wing size and shape in Eudamina skippers

Ancestral state reconstruction revealed the independent evolution of hindwing tails and dorsal iridescence in multiple lineages, with repeated losses and gains across genera (e.g., *Cecropterus* and *Telegonus*). As both traits are more common in tropical environments than in the subtropics, a possible explanation is that these evolve under high predation pressure and diverse types of predators (Sherry et al., 2020). To test this hypothesis, accurate data on species distributions, of both predators and prey, are needed to evaluate patterns of co-occurrences.

The macroevolutionary dynamics of all morphometric traits were best explained by Ornstein-Uhlenbeck models, pointing to evolution under constraints rather than unconstrained drift or early burst dynamics. In every case OU outperformed both Brownian motion (BM) and early burst (EB) alternatives by margins of ΔAICc ≫ 50. This consistency indicates that each trait has evolved toward and is maintained around fitness optima, reflecting stabilizing selection across Eudamina. For hindwing tail length in particular, the OU fit implies two discrete peaks (tail absence or presence of long tails) and the near absence of intermediates, as shown on Figure 3 or 5. Together, these results show that rather than drifting freely or radiating explosively, the morphologies and flight proxies have been repeatedly pulled back toward alternative states throughout the evolutionary history of this group. Future studies could compare these traits states to pinpoint the selective pressures driving convergent evolution in independent lineages.

In conclusion, our results show that convergent traits in Eudamina skippers are evolutionarily correlated with flight proxies indicative of high escape ability. Dorsal iridescence is associated with high wing loading and aspect ratio, indicative of a fast flight, in agreement with the hypothesis of Janzen et al., (2009). Creamy bands in turn, were not associated with any recurrent wing morphology and seem to evolve neutrally in Eudamina. Tails are associated with intermediate body size and reduced wing loading, indicative of a lower speed, but also with a forewing shape that might improve flight agility. They might thus directly contribute to escape ability. Alternatively, they might deflect predator attacks, thereby imposing recurrent constraints on forewing shape, suggesting that both wings do not evolve independently. Combined with the strong support for OU models of evolution, these results suggests that selective pressures related to predation have shaped Eudamina skippers wing convergences.

## Acknowledgment

Firstly, we would like to thank all museum curators who opened their collections for our study: Théo Léger (MfN, Berlin), Blanca Huertas (NHM, London) and Gerardo Lamas (UNMSM, Lima). Additionally, we would like to thank Rayner Nuñez for photos of *Chioides marmorosa*, Alena Sucháčková for comments during review and help during figure preparation, as well as Prapti Gohil and Ema Ritterová for their help during image selection and measuring of wing characteristics.

## Authors contributions

**DL** – study planning, data collection and evaluation, writing original draft, review and editing; **VD** – review and editing; **PMM** – study planning, funding acquisition, review and editing.

## Data availability

The data that support the findings of this study are available from the corresponding author upon reasonable request and will be archived on Zenodo after acceptance.

## References

Adams, D. C., & Otárola-Castillo, E. (2013). geomorph: An r package for the collection and analysis of geometric morphometric shape data. Methods in Ecology and Evolution, 4(4), 393–399. 10.1111/2041-210X.12035

Bertrand, C., Janzen, D., Hallwachs, W., Burns, J., Gibson, J., Shokralla, S., & Hajibabaei, M. (2014). Mitochondrial and nuclear phylogenetic analysis with Sanger and next-generation sequencing shows that, in Área de Conservación Guanacaste, northwestern Costa Rica, the skipper butterfly named Urbanus belli (family Hesperiidae) comprises three morphologically cryptic species. BMC Evolutionary Biology, 14, 153. 10.1186/1471-2148-14-153

Berwaerts, K., Van Dyck, H., & Aerts, P. (2002). Does Flight Morphology Relate to Flight Performance? An Experimental Test with the Butterfly Pararge aegeria. Functional Ecology, 16(4), 484–491.

Betts, C. R., & Wootton, R. J. (1988). Wing Shape and Flight Behaviour in Butterflies (Lepidoptera: Papilionoidea and Hesperioidea): A Preliminary Analysis. Journal of Experimental Biology, 138(1), 271–288. 10.1242/jeb.138.1.271

Borics, G., Várbíró, G., Falucskai, J., Végvári, Z., T-Krasznai, E., Görgényi, J., B-Béres, V., & Lerf, V. (2023). A two-dimensional morphospace for cyanobacteria and microalgae: Morphological diversity, evolutionary relatedness, and size constraints. Freshwater Biology, 68(1), 115–126. 10.1111/fwb.14013

Bouckaert, R., Heled, J., Kühnert, D., Vaughan, T., Wu, C.-H., Xie, D., Suchard, M. A., Rambaut, A., & Drummond, A. J. (2014). BEAST 2: A Software Platform for Bayesian Evolutionary Analysis. PLOS Computational Biology, 10(4), e1003537. 10.1371/journal.pcbi.1003537

Bürkner, P.-C. (2017). brms: An R package for Bayesian multilevel models using Stan. Journal of Statistical Software, 80, 1–28.

Cespedes, A., Penz, C. M., & DeVries, P. J. (2015). Cruising the rain forest floor: Butterfly wing shape evolution and gliding in ground effect. Journal of Animal Ecology, 84(3), 808–816. 10.1111/1365-2656.12325

Chotard, A., Ledamoisel, J., Decamps, T., Herrel, A., Chaine, A. S., Llaurens, V., & Debat, V. (2022). Evidence of attack deflection suggests adaptive evolution of wing tails in butterflies. Proceedings of the Royal Society B: Biological Sciences, 289(1975), 20220562. 10.1098/rspb.2022.0562

Clavel, J., Escarguel, G., & Merceron, G. (2015). mvmorph: An r package for fitting multivariate evolutionary models to morphometric data. Methods in Ecology and Evolution, 6(11), 1311– 1319. 10.1111/2041-210X.12420

Darwin, C. (1859). On the origin of species by means of natural selection, or, The preservation of favoured races in the struggle for life. J. Murray. https://www.loc.gov/resource/rbctos.2017gen17473/

DeVries, P. J., Penz, C. M., & Hill, R. I. (2010). Vertical distribution, flight behaviour and evolution of wing morphology in Morpho butterflies. Journal of Animal Ecology, 79(5), 1077–1085. 10.1111/j.1365-2656.2010.01710.x

Dudley, R. (2002). The Biomechanics of Insect Flight: Form, Function, Evolution. Princeton University Press.

Galis, F., Metz, J. A. J., & Alphen, J. J. M. van. (2018). Development and Evolutionary Constraints in Animals. Annual Review of Ecology, Evolution, and Systematics, 49(Volume 49, 2018), 499–522. 10.1146/annurev-ecolsys-110617-062339

Hamilton, C. A., Winiger, N., Rubin, J. J., Breinholt, J., Rougerie, R., Kitching, I. J., Barber, J. R., & Kawahara, A. Y. (2022). Hidden Phylogenomic Signal Helps Elucidate Arsenurine Silkmoth Phylogeny and the Evolution of Body Size and Wing Shape Trade-Offs. Systematic Biology, 71(4), 859–874. 10.1093/sysbio/syab090

Hendrick, L. K., Somjee, U., Rubin, J. J., & Kawahara, A. Y. (2022). A Review of False Heads in Lycaenid Butterflies. The Journal of the Lepidopterists’ Society, 76(2), 140–148. 10.18473/lepi.76i2.a6

Henriques, N. R., Lourenço, G. M., Diniz, É. S., & Cornelissen, T. (2022). Is elevation a strong environmental filter? Combining taxonomy, functional traits and phylogeny of butterflies in a tropical mountain. Ecological Entomology, 47(4), 613–625. 10.1111/een.13145

Holling, C. S. (1965). The Functional Response of Predators to Prey Density and its Role in Mimicry and Population Regulation. The Memoirs of the Entomological Society of Canada, 97(S45), 5–60. 10.4039/entm9745fv

Jahant-Miller, C., Miller, R., & Parry, D. (2022). Size-dependent flight capacity and propensity in a range-expanding invasive insect. Insect Science, 29(3), 879–888. 10.1111/1744-7917.12950

Jantzen, B., & Eisner, T. (2008). Hindwings are unnecessary for flight but essential for execution of normal evasive flight in Lepidoptera. Proceedings of the National Academy of Sciences, 105(43), 16636–16640. 10.1073/pnas.0807223105

Janzen, D. H., Hallwachs, W., Blandin, P., Burns, J. M., Cadiou, J.-M., Chacon, I., Dapkey, T., Deans, A. R., Epstein, M. E., Espinoza, B., Franclemont, J. G., Haber, W. A., Hajibabaei, M., Hall, J. P. W., Hebert, P. D. N., Gauld, I. D., Harvey, D. J., Hausmann, A., Kitching, I. J.,…Wilson, J. J. (2009). Integration of DNA barcoding into an ongoing inventory of complex tropical biodiversity. Molecular Ecology Resources, 9(s1), 1–26. 10.1111/j.1755-0998.2009.02628.x

Jyothi, P., Aralimarad, P., Wali, V., Dave, S., Bheemanna, M., Ashoka, J., Shivayogiyappa, P., Lim, K. S., Chapman, J. W., & Sane, S. P. (2021). Evidence for facultative migratory flight behavior in Helicoverpa armigera (Noctuidae: Lepidoptera) in India. PLOS ONE, 16(1), e0245665. 10.1371/journal.pone.0245665

Kang, C., Zahiri, R., & Sherratt, T. N. (2017). Body size affects the evolution of hidden colour signals in moths. Proceedings of the Royal Society B: Biological Sciences, 284(1861), 20171287. 10.1098/rspb.2017.1287

Kawahara, A. Y., Storer, C., Carvalho, A. P. S., Plotkin, D. M., Condamine, F. L., Braga, M. P., Ellis, E. A., St Laurent, R. A., Li, X., Barve, V., Cai, L., Earl, C., Frandsen, P. B., Owens, H. L., Valencia-Montoya, W. A., Aduse-Poku, K., Toussaint, E. F. A., Dexter, K. M., Doleck, T.,…Lohman, D. J. (2023). A global phylogeny of butterflies reveals their evolutionary history, ancestral hosts and biogeographic origins. Nature Ecology & Evolution, 7(6), Article 6. 10.1038/s41559-023-02041-9

Kleckova, I., Linke, D., Rezende, F. D. M., Rauscher, L., Le Roy, C., & Matos-Maraví, P. (2024). Flight behaviour diverges more between seasonal forms than between species in Pieris butterflies. Ecology and Evolution, 14(7), e70012. 10.1002/ece3.70012

Le Roy, C., Debat, V., & Llaurens, V. (2019). Adaptive evolution of butterfly wing shape: From morphology to behaviour. Biological Reviews, 94(4), 1261–1281. 10.1111/brv.12500

Li, W., Cong, Q., Shen, J., Zhang, J., Hallwachs, W., Janzen, D. H., & Grishin, N. V. (2019). Genomes of skipper butterflies reveal extensive convergence of wing patterns. Proceedings of the National Academy of Sciences, 116(13), 6232–6237. 10.1073/pnas.1821304116

Linke, D., Elias, M., Klečková, I., Mappes, J., & Matos-Maraví, P. (2022). Shape of Evasive Prey Can Be an Important Cue That Triggers Learning in Avian Predators. Frontiers in Ecology and Evolution, 10. 10.3389/fevo.2022.910695

Linke, D., Hernandez Mejia, J., N. P. Eche Navarro, V., Salinas Sánchez, L., de Gusmão Ribeiro, P., Elias, M., & Matos-Maraví, P. (2024). Reduced palatability, fast flight, and tails: Decoding the defence arsenal of Eudaminae skipper butterflies in a Neotropical locality. Journal of Evolutionary Biology, voae091. 10.1093/jeb/voae091

Loeffler-Henry, K., & Sherratt, T. N. (2024). Selection for evasive mimicry imposed by an arthropod predator. Biology Letters, 20(1), 20230461. 10.1098/rsbl.2023.0461

Mitteroecker, P., & Gunz, P. (2009). Advances in Geometric Morphometrics. Evolutionary Biology, 36(2), 235–247. 10.1007/s11692-009-9055-x

Morrone, J. J. (2014). Cladistic biogeography of the Neotropical region: Identifying the main events in the diversification of the terrestrial biota. Cladistics, 30(2), 202–214. 10.1111/cla.12039

Murali, G. (2018). Now you see me, now you don’t: Dynamic flash coloration as an antipredator strategy in motion. Animal Behaviour, 142, 207–220. 10.1016/j.anbehav.2018.06.017

Owens, H. L., Lewis, D. S., Condamine, F. L., Kawahara, A. Y., & Guralnick, R. P. (2020). Comparative Phylogenetics of Papilio Butterfly Wing Shape and Size Demonstrates Independent Hindwing and Forewing Evolution. Systematic Biology, 69(5), 813–819. 10.1093/sysbio/syaa029

Páez, E., Valkonen, J. K., Willmott, K. R., Matos-Maraví, P., Elias, M., & Mappes, J. (2021). Hard to catch: Experimental evidence supports evasive mimicry. Proceedings. Biological Sciences, 288(1946), 20203052. 10.1098/rspb.2020.3052

Paradis, E., & Schliep, K. (2019). ape 5.0: An environment for modern phylogenetics and evolutionary analyses in R. Bioinformatics, 35(3), 526–528.

Park, H., Bae, K., Lee, B., Jeon, W.-P., & Choi, H. (2010). Aerodynamic Performance of a Gliding Swallowtail Butterfly Wing Model. Experimental Mechanics, 50(9), 1313–1321. 10.1007/s11340-009-9330-x

Pinheiro, C. E. G., & Freitas, A. V. L. (2014). Some Possible Cases of Escape Mimicry in Neotropical Butterflies. Neotropical Entomology, 43(5), 393–398. 10.1007/s13744-014-0240-y

Qing, Y., & Zhi-tao, Z. (2001). Analysis of Wing-Shape Characteritics of Migratory Lepidopterous Insects. Insect Science, 8(2), 183–192. 10.1111/j.1744-7917.2001.tb00484.x

Revell, L. J. (2012). phytools: An R package for phylogenetic comparative biology (and other things). Methods in Ecology and Evolution, 3(2), 217–223. 10.1111/j.2041-210X.2011.00169.x

Ribeiro, P. G., Torres Jiménez, M. F., Andermann, T., Antonelli, A., Bacon, C. D., & Matos-Maraví, P. (2021). A bioinformatic platform to integrate target capture and whole genome sequences of various read depths for phylogenomics. Molecular Ecology, 30(23), 6021–6035. 10.1111/mec.16240

Robbins, R. K. (1981). The ‘False Head’ Hypothesis: Predation and Wing Pattern Variation of Lycaenid Butterflies. The American Naturalist, 118(5), 770–775. 10.1086/283868

Rohlf, F. J. (2005). tpsDig: Digitize landmarks and outlines. Version 2.05 [computer program]. Department of Ecology and Evolution, State University of New York at Stony Brook, Stony Brook.

Rossato, D. O., Kaminski, L. A., Iserhard, C. A., & Duarte, L. (2018). Chapter Two - More Than Colours: An Eco-Evolutionary Framework for Wing Shape Diversity in Butterflies. In R. H. ffrench-Constant (Ed.), Advances in Insect Physiology (Vol. 54, pp. 55–84). Academic Press. 10.1016/bs.aiip.2017.11.003

Rubin, J. J., Hamilton, C. A., McClure, C. J. W., Chadwell, B. A., Kawahara, A. Y., & Barber, J. R. (2018). The evolution of anti-bat sensory illusions in moths. Science Advances, 4(7), eaar7428. 10.1126/sciadv.aar7428

Sherry, T. W., Kent, C. M., Sánchez, N. V., & Şekercioğlu, Ç. H. (2020). Insectivorous birds in the Neotropics: Ecological radiations, specialization, and coexistence in species-rich communities. The Auk, 137(4), ukaa049. 10.1093/auk/ukaa049

Silvasti, S., Kemp, D. J., White, T. E., Nokelainen, O., Valkonen, J., & Mappes, J. (2024). The flashy escape: Support for dynamic flash coloration as anti-predator defence. Biology Letters, 20(7), 20240303. 10.1098/rsbl.2024.0303

Tendler, A., Mayo, A., & Alon, U. (2015). Evolutionary tradeoffs, Pareto optimality and the morphology of ammonite shells. BMC Systems Biology, 9(1), 12. 10.1186/s12918-015-0149-z

van Someren, V. G. L., & Jackson, T. H. E. (1959). Some comments on protective resemblance amongst African Lepidoptera (Rhopalocera). The Lepidopterist Society, 13(3), 121–150.

Vieira-Silva, A., Evora, G. B., Freitas, A. V. L., & Oliveira, P. S. (2024). The Relevance of Flash Coloration Against Avian Predation in a Morpho Butterfly: A Field Experiment in a Tropical Rainforest. Ethology, 130(12), e13517. 10.1111/eth.13517

Wickham, H. (2016). Ggplot2. Springer International Publishing. 10.1007/978-3-319-24277-4

Zhang, J., Cong, Q., & Grishin, N. V. (2023). Thirteen new species of butterflies (Lepidoptera: Hesperiidae) from Texas. Insecta Mundi, 2023, 0969.

Zhang, J., Cong, Q., Shen, J., Brockmann, E., & Grishin, N. V. (2019). Genomes reveal drastic and recurrent phenotypic divergence in firetip skipper butterflies (Hesperiidae: Pyrrhopyginae). Proceedings. Biological Sciences, 286(1903), 20190609. 10.1098/rspb.2019.0609

