## Appendix for "Convergent evolution of skipper wings assessed through anti-predator defences, flight proxies and geographical distribution"

**Appendix S1:** Table for genetic data

**Table S1:** Table indicating collection locality (State/Province, Country), storage location (i.e., museum or university as well as accession codes for online repositories. For species marked with HOLDER no genetic sequences were found, these species were inserted at random into their corresponding genus.

| **Species** | **Code** | **Collection** | **Specimen storage** | **Online repository** | **Accession number** |
| --- | --- | --- | --- | --- | --- |
| Outgroup *Aguna spicata* | PM0300 | Madre de Dios, Peru | Museo de Historia Natural, Lima | BOLD | LEPPE099-22 |
| *Astraptes aulus* | BN000149 |  | E. Tuissant | not submitted | not submitted |
| *Astraptes enotrus* | BCI91003 | Provincia de Panama, Panama | Collection České Budějovice | not submitted | not submitted |
| *Astraptes erycina* | HOLDER |  |  |  |  |
| *Astraptes halesius* | HOLDER |  |  |  |  |
| *Astraptes janeira* | DQ291884 | Alajuela, Costa Rica |  | NCBI | DQ291884 |
| *Astraptes mabillei* | MH357768 | Cusco, Peru |  | NCBI | MH357768 |
| *Autochton bipunctatus* | NVG-5748 | Madre de Dios, Peru | Collection České Budějovice | not submitted | not submitted |
| *Autochton caballo* | OP762106 | Texas, USA |  | NCBI | OP762106 |
| *Autochton integrifascia* | MH357766 | Minas Gerais, Brazil |  | NCBI | MH357766 |
| *Autochton itylus* | PM0117 | Pasco, Peru | Collection České Budějovice | BOLD | LEPPE024-22 |
| *Autochton neis* | BN000160 |  | E. Tuissant | not submitted | not submitted |
| *Autochton oryx* | NVG-5693 | Sucumbios, Ecuador | National Museum of Natural History, Smithsonian Institution, Washington, DC | NCBI | SRR7174543 |
| *Autochton potrillo* | HM885925 | Guanacaste, Costa Rica |  | NCBI | HM885925 |
| *Autochton reducta* | HOLDER |  |  |  |  |
| *Autochton reflexus* | NVG-15111F10 | Santa Catarina, Brazil | American Museum of Natural History, New York | NCBI | SRR7174388 |
| *Autochton sulfureolus* | MH357764 | Rio de Janeiro, Brazil |  | NCBI | MH357764 |
| *Cecropters markwalkeri* | OQ311409_1 | Sonora, Mexico |  | NCBI | OQ311409_1 |
| *Cecropterus acanthopoda* | NVG-15101A07 | Rio Grande do Sul, Brazil | National Museum of Natural History, Smithsonian Institution, Washington, DC | NCBI | SRR7174384 |
| *Cecropterus albimargo* | LEP41551 |  | E. Tuissant | not submitted | not submitted |
| *Cecropterus albociliatus* | NVG-5749 | Guanacaste, Costa Rica | National Museum of Natural History, Smithsonian Institution, Washington, DC | NCBI | SRR7174578 |
| *Cecropterus athesis* | HOLDER |  |  |  |  |
| *Cecropterus barra* | HOLDER |  |  |  |  |
| *Cecropterus bathyllus* | NVG-4539 | Oklahoma, USA |  | NCBI | SRR7174390 |
| *Cecropterus carmelita* | HOLDER |  |  |  |  |
| *Cecropterus casica* | NVG-8269 | Arizona, USA |  | NCBI | SRR7174392 |
| *Cecropterus cincta* | MH357755_1 | Texas, USA |  | NCBI | MH357755_1 |
| *Cecropterus confusis* | NVG-4185 | Texas, USA |  | NCBI | SRR7174394 |
| *Cecropterus coyote* | NVG-3830 | Texas, USA |  | NCBI | SRR7174573 |
| *Cecropterus cramptoni* | JN270038_1 | Dominican Republic |  | NCBI | JN270038_1 |
| *Cecropterus diversus* | MW807655_1 | California, USA |  | NCBI | MW807655_1 |
| *Cecropterus dobra* | LEP31595 |  | E. Tuissant | not submitted | not submitted |
| *Cecropterus dorantes* | PM-UN-11 | Cienfuegos, Cuba | SNSB, Zoologische Staatssammlung, München | not submitted | not submitted |
| *Cecropterus doryssus* | BCI142105 | Provincia de Panama, Panama | Collection České Budějovice | not submitted | not submitted |
| *Cecropterus drusius* | LNAUW318-17 | Arizona, USA | National Museum of Natural History, Smithsonian Institution, Washington, DC | BOLD | LNAUW318-17 |
| *Cecropterus egregius* | BCI150806 | Provincia de Panama, Panama | Collection České Budějovice | not submitted | not submitted |
| *Cecropterus evenus* | NVG-5072 | Minas Gerais, Brazil | National Museum of Natural History, Smithsonian Institution, Washington, DC | NCBI | SRR7174399 |
| *Cecropterus jalapus* | BLPAA3239-17 | Guanacaste, Costa Rica | University of Pennsylvania | BOLD | BLPAA3239-17 |
| *Cecropterus longipennis* | BCI91297 | Provincia de Panama, Panama | Smithsonian Tropical Research Institute, Panama | not submitted | not submitted |
| *Cecropterus lyciades* | NVG-3311 | Texas, USA |  | not submitted | not submitted |
| *Cecropterus mexicana* | MH357757_1 | Oaxaca, Mexico |  | NCBI | MH357757_1 |
| *Cecropterus nevada* | ABLCV133-09 | California, USA | Colorado State University | BOLD | ABLCV133-09 |
| *Cecropterus nigrociliata* | BN003357 |  | E. Tuissant | not submitted | not submitted |
| *Cecropterus obscurus* | NVG-22044E09 |  |  | not submitted | not submitted |
| *Cecropterus palliolum* | NVG-14104A05 | Limon, Costa Rica | National Museum of Natural History, Smithsonian Institution, Washington, DC | NCBI | SRR7174576 |
| *Cecropterus phalaecus* | NVG-14103G09 | San Luis Potosi, Mexico | National Museum of Natural History, Smithsonian Institution, Washington, DC | NCBI | SRR7174581 |
| *Cecropterus pseudocellus* | NVG-14061G02 | Durango, Mexico | National Museum of Natural History, Smithsonian Institution, Washington, DC | NCBI | SRR7174395 |
| *Cecropterus pylades* | LEP39973 |  | E. Tuissant | not submitted | not submitted |
| *Cecropterus reductus* | MH357760_1 | Guyana |  | NCBI | MH357760_1 |
| *Cecropterus rica* | HOLDER |  |  |  |  |
| *Cecropterus rinta* | NVG-15026C12 | Salta, Argentina | McGuire Center for Lepidoptera and Biodiversity, Gainesville | NCBI | SRR7174396 |
| *Cecropterus tehuacana* | 11-BOA-15609H02 | Coahuila, Mexico | National Museum of Natural History, Smithsonian Institution, Washington, DC | NCBI | SRR7174393 |
| *Cecropterus trebia* | MH357761_1 | Porto Cabello, Venezuela |  | NCBI | MH357761_1 |
| *Cecropterus vectilucis* | MH357756_1 | Chiriqui, Panama |  | NCBI | MH357756_1 |
| *Cecropterus virescence* | MH357759_1 | Guyana |  | NCBI | MH357759_1 |
| *Cecropterus zarex* | BLPAA3317-17 | Guanacaste, Costa Rica | University of Pennsylvania | BOLD | BLPAA3317-17 |
| *Cecroptrus takuta* | NVG-15093A09 | Rondonia, Brazil | McGuire Center for Lepidoptera and Biodiversity, Gainesville | NCBI | SRR7174575 |
| *Chioides albofasciatus* | NVG-4502 | Texas, USA |  | NCBI | SRR7174540 |
| *Chioides catillus* | 151-ADW | Guanacaste, Costa Rica |  | NCBI | KY019706 |
| *Chioides churchi* | JN269969 | Jamaica |  | NCBI | JN269969 |
| *Chioides cinereus* | HOLDER |  |  |  |  |
| *Chioides concinnus* | HOLDER |  |  |  |  |
| *Chioides iverna* | HOLDER |  |  |  |  |
| *Chioides ixion* | OP586833 | La Altagracia, Dom. Rep. |  | NCBI | OP586833 |
| *Chioides marmorosa* | PM-UN-10 | Artemisa, Cuba | SNSB, Zoologische Staatssammlung München | not submitted | not submitted |
| *Chioides vintra* | NVG-5053 | Lowmans, St. Vincent | National Museum of Natural History, Smithsonian Institution, Washington, DC | NCBI | SRR7174541 |
| *Chioides zilpa* | DQ292134 | Guanacaste, Costa Rica |  | NCBI | DQ292134 |
| *Epargyerus enispe* | HOLDER |  |  |  |  |
| *Epargyreus antaeus* | HOLDER |  |  |  |  |
| *Epargyreus aspina* | HOLDER |  |  |  |  |
| *Epargyreus barisses* | DL1260 | Cusco, Peru | Museo de Historia Natural, Lima |  | not submitted |
| *Epargyreus brodkorbi* | HOLDER |  |  |  |  |
| *Epargyreus clarus* | NVG-4192 | Texas, USA | Nick Grishin? | NCBI | SRR7174549 |
| *Epargyreus clavicornis* | DL1261 | Madre de Dios, Peru | Museo de Historia Natural, Lima | not submitted | not submitted |
| *Epargyreus cruza* | HM885893 | Alajuela, Costa Rica |  | NCBI | HM885893 |
| *Epargyreus deleoni* | GU658305 | Mexico |  | NCBI | GU658305 |
| *Epargyreus dicta* | DL1262 | Cusco, Peru | Museo de Historia Natural, Lima | not submitted | not submitted |
| *Epargyreus exadeus* | MF545889 | Misiones, Argentina |  | NCBI | MF545889 |
| *Epargyreus fractigutta* | OP762107 | Texas, USA |  | NCBI | OP762107 |
| *Epargyreus gaumeri* | HOLDER |  |  |  |  |
| *Epargyreus huachuca* | HOLDER |  |  |  |  |
| *Epargyreus nutra* | HOLDER |  |  |  |  |
| *Epargyreus orizaba* | HOLDER |  |  |  |  |
| *Epargyreus pseudexadeus* | HOLDER |  |  |  |  |
| *Epargyreus socus* | MH357769 | Guyana |  | NCBI | MH357769 |
| *Epargyreus spanna* | HOLDER |  |  |  |  |
| *Epargyreus spina* | GU658356 | Mexico |  | NCBI | GU658356 |
| *Epargyreus spinosa* | HOLDER |  |  |  |  |
| *Epargyreus spinta* | HOLDER |  |  |  |  |
| *Epargyreus tmolis* | MF546835 | Entre Rios, Argentina |  | NCBI | MF546835 |
| *Epargyreus windi* | HOLDER |  |  |  |  |
| *Epargyreus zestos* | JN277829 | Puerto Rico |  | NCBI | JN277829 |
| *Narcosius aulina* | HOLDER |  |  |  |  |
| *Narcosius colossus* | JF762384 | Alajuela, Costa Rica |  | NCBI | JF762384 |
| *Narcosius dosula* | HOLDER |  |  |  |  |
| *Narcosius granadensis* | HOLDER |  |  |  |  |
| *Narcosius helen* | NVG-15081D11 | Veracruz, Mexico |  | NCBI | SRR7174546 |
| *Narcosius hercules* | DL1265 | Madre de Dios, Peru | Museo de Historia Natural, Lima | not submitted | not submitted |
| *Narcosius mura* | DL1266 | Madre de Dios, Peru | Museo de Historia Natural, Lima | not submitted | not submitted |
| *Narcosius narcosius* | HQ567237 | Brazil |  | NCBI | HQ567237 |
| *Narcosius nazaeus* | PM0309 | Madre de Dios, Peru | Museo de Historia Natural, Lima | BOLD | LEPPE106-22 |
| *Narcosius odysseus* | HOLDER |  |  |  |  |
| *Narcosius parisi* | HOLDER |  |  |  |  |
| *Narcosius pseudomura* | DL1267 | Madre de Dios, Peru | Museo de Historia Natural, Lima | not submitted | not submitted |
| *Narcosius samson* | BCI112538 | Panama Oeste, Panama |  | NCBI | BCI112538 |
| *Narcosius steinhauseri* | HOLDER |  |  |  |  |
| *Proteides maysi* | OP587044 | Holguín, Cuba |  | NCBI | OP587044 |
| *Proteides mercurius* | ZSM-91542 | Holguín, Cuba | SNSB, Zoologische Staatssammlung, München | not submitted | not submitted |
| *Spathlepia clonius* | NVG-3835 | Texas, USA |  | NCBI | SRR7174548 |
| *Spicauda ambiguus* | HOLDER |  |  |  |  |
| *Spicauda atelis* | OP984702 | Texas, USA |  | NCBI | OP984702 |
| *Spicauda cindra* | PM0107 | Pasco, Peru | Collection České Budějovice | BOLD | LEPPE019-22 |
| *Spicauda procne* | NVG-3754 | Texas, USA |  | NCBI | SRR7174580 |
| *Spicauda simplicius* | PM0123 | Pasco, Peru | Collection České Budějovice | BOLD | LEPPE027-22 |
| *Spicauda tanna* | PM0069 | Junin, Peru | Collection České Budějovice | BOLD | LEPPE007-22 |
| *Spicauda teleus* | BN000868 |  | E. Tuissant | not submitted | not submitted |
| *Spicauda zagorus* | BN002827 |  | E. Tuissant | not submitted | not submitted |
| *Spicauda zalanthus* | MF545923 | Misiones, Argentina |  | NCBI | MF545923 |
| *Telegonus alardus* | BN003359 |  | E. Tuissant | not submitted | not submitted |
| *Telegonus alector* | NVG-5076 | Alajuela, Costa Rica | National Museum of Natural History, Smithsonian Institution, Washington, DC | NCBI | SRR7174510 |
| *Telegonus anaphus* | BCI90750 | Provincia de Panama, Panama | Collection České Budějovice | not submitted | not submitted |
| *Telegonus anausis* | BLPAA11588-18 | Alajuela, Costa Rica | University of Pennsylvania | BOLD | BLPAA11588-18 |
| *Telegonus apastus* | NVG-5078 | Alajuela, Costa Rica | National Museum of Natural History, Smithsonian Institution, Washington, DC | NCBI | SRR7174511 |
| *Telegonus azul* | BCI150133 | Provincia de Panama, Panama | Smithsonian Tropical Research Institute, Panama | not submitted | not submitted |
| *Telegonus bifascia* | HOLDER |  |  |  |  |
| *Telegonus brevicauda* | NVG-17096A09 | Alajuela, Costa Rica | National Museum of Natural History, Smithsonian Institution, Washington, DC | NCBI | SRR7174348 |
| *Telegonus cassander* | PM-UN-15 | Granma, Cuba | SNSB, Zoologische Staatssammlung, München | not submitted | not submitted |
| *Telegonus cassius* | MH357777 | Puntarenes, Costa Rica |  | NCBI | MH357777 |
| *Telegonus catemacoensis* | BLPAA18927-20 | Guanacaste, Costa Rica | University of Pennsylvania | BOLD | BLPAA18927-20 |
| *Telegonus cellus* | HQ561161 | USA |  | NCBI | HQ561161 |
| *Telegonus chalco* | BCI86454 | Provincia de Panama, Panama | Smithsonian Tropical Research Institute, Panama | not submitted | not submitted |
| *Telegonus chiriquensis* | DL1268 | Cusco, Peru | Museo de Historia Natural, Lima | not submitted | not submitted |
| *Telegonus christyi* | KY659596 | Le Vega, Dom. Rep. |  | NCBI | KY659596 |
| *Telegonus cretatus* | HM394667 | Bolivia |  | NCBI | HM394667 |
| *Telegonus cretellus* | MH357778 | Trelawny, Jamaica |  | NCBI | MH357778 |
| *Telegonus creteus* | 00-BCI-2545 | Panama | Smithsonian Tropical Research Institute, Panama | not submitted | not submitted |
| *Telegonus elorus* | PM-UN-24 | Minas Gerais, Brazil | Museu de História Natural, UNICAMP, Brazil | not submitted | not submitted |
| *Telegonus fulgerator* | BCI135008 | Provincia de Panama, Panama | Collection České Budějovice | not submitted | not submitted |
| *Telegonus fulgor* | HQ567093 | Argentina |  | NCBI | HQ567093 |
| *Telegonus fulminator* | OP762102 | Suriname |  | NCBI | OP762102 |
| *Telegonus galesus* | MH357776 | Cusco, Peru |  | NCBI | MH357776 |
| *Telegonus habana* | PM-UN-13 | Granma, Cuba | SNSB, Zoologische Staatssammlung, München | not submitted | not submitted |
| *Telegonus heriul* | JN270044 | Dominican Republic |  | NCBI | JN270044 |
| *Telegonus hyster* | NVG-5676 | Guerrero, Mexico | National Museum of Natural History, Smithsonian Institution, Washington, DC | NCBI | SRR7174508 |
| *Telegonus latimargo* | HOLDER |  |  |  |  |
| *Telegonus misitra* | OP762104 | Mexico |  | NCBI | OP762104 |
| *Telegonus naxos* | PM-UN-25 | Minas Gerais, Brazil | Museu de História Natural, UNICAMP, Brazil | not submitted | not submitted |
| *Telegonus siermadror* | MH357773 | Nuevo Leon, Mexico |  | NCBI | MH357773 |
| *Telegonus subflavus* | NVG-15096B05 |  |  | Zhan et al. 2022 | 10.5281/zenodo.6392056 |
| *Telegonus talus* | ZSM-91566 | Holguín, Cuba | Feliberto Bermudez Research Collection | NCBI | OP587187 |
| *Telegonus tinda* | HOLDER |  |  |  |  |
| *Telegonus tsongae* | OP762105 | Texas, USA |  | NCBI | OP762105 |
| *Telegonus weymeri* | HOLDER |  |  |  |  |
| *Telegonus xagua* | ZSM-92857 | Guantanamo, Cuba | Museo Nacional de Historia Natural Cuba, MNHNC | not submitted | not submitted |
| *Urbanus alva* | BLPAA3443-17 | Alajuela, Costa Rica | University of Pennsylvania | BOLD | BLPAA3443-17 |
| *Urbanus belli* | BCI32672 | Provincia de Panama, Panama | Collection České Budějovice | not submitted | not submitted |
| *Urbanus bernikerni* | BLPAA3451-17 | Guanacaste, Costa Rica | University of Pennsylvania | BOLD | BLPAA3451-17 |
| *Urbanus dubius* | HOLDER |  |  |  |  |
| *Urbanus elmina* | PM0086 | Pasco, Peru | Collection České Budějovice | BOLD | LEPPE011-22 |
| *Urbanus esma* | HM416551 | Panama |  | NCBI | HM416551 |
| *Urbanus esmeraldus* | NVG-5691 | Alajuela, Costa Rica | National Museum of Natural History, Smithsonian Institution, Washington, DC | NCBI | SRR7174353 |
| *Urbanus esta* | BLPAA3477-17 | Guanacaste, Costa Rica | University of Pennsylvania | BOLD | BLPAA3477-17 |
| *Urbanus evona* | NVG-15103F04 | Guanacaste, Costa Rica | National Museum of Natural History, Smithsonian Institution, Washington, DC | NCBI | SRR7174354 |
| *Urbanus huancavillcas* | HOLDER |  |  |  |  |
| *Urbanus longicaudus* | HOLDER |  |  |  |  |
| *Urbanus magnus* | HOLDER |  |  |  |  |
| *Urbanus megalurus* | NVG-14082H06 | Chiapas, Mexico | American Museum of Natural History, New York | NCBI | SRR7174350 |
| *Urbanus oplerorum* | OP762100 | Texas, USA |  | NCBI | OP762100 |
| *Urbanus parvus* | MH357771 | Guyana |  | NCBI | MH357771 |
| *Urbanus prodicus* | MH357770 | Nuevo Leon, Mexico |  | NCBI | MH357770 |
| *Urbanus pronta* | DQ293769 | Guanacaste, Costa Rica |  | NCBI | DQ293769 |
| *Urbanus pronus* | 99-BCI-1601 | Provincia de Panama, Panama | Smithsonian Tropical Research Institute, Panama | not submitted | not submitted |
| *Urbanus proteus* | NVG-4894 | Florida, USA |  | NCBI | SRR7174352 |
| *Urbanus rickardi* | OP762099 | Texas, USA |  | NCBI | OP762099 |
| *Urbanus segnestami* | NVG-15103E08 | Alajuela, Costa Rica | National Museum of Natural History, Smithsonian Institution, Washington, DC | NCBI | SRR7174349 |
| *Urbanus tucuti* | NVG-2932 | Veracruz, Mexico |  | NCBI | SRR7174347 |
| *Urbanus velinus* | HSPED947-12 | Pastaza, Ecuador | Research Collectio of Ernst Brockmann | BOLD | HSPED947-12 |
| *Urbanus villus* | HOLDER |  |  |  |  |
| *Urbanus viridis* | HOLDER |  |  |  |  |
| *Urbanus viterboana* | DL1269 | Cusco, Peru | Museo de Historia Natural, Lima | not submitted | not submitted |

**Appendix S2:** Full consensus tree

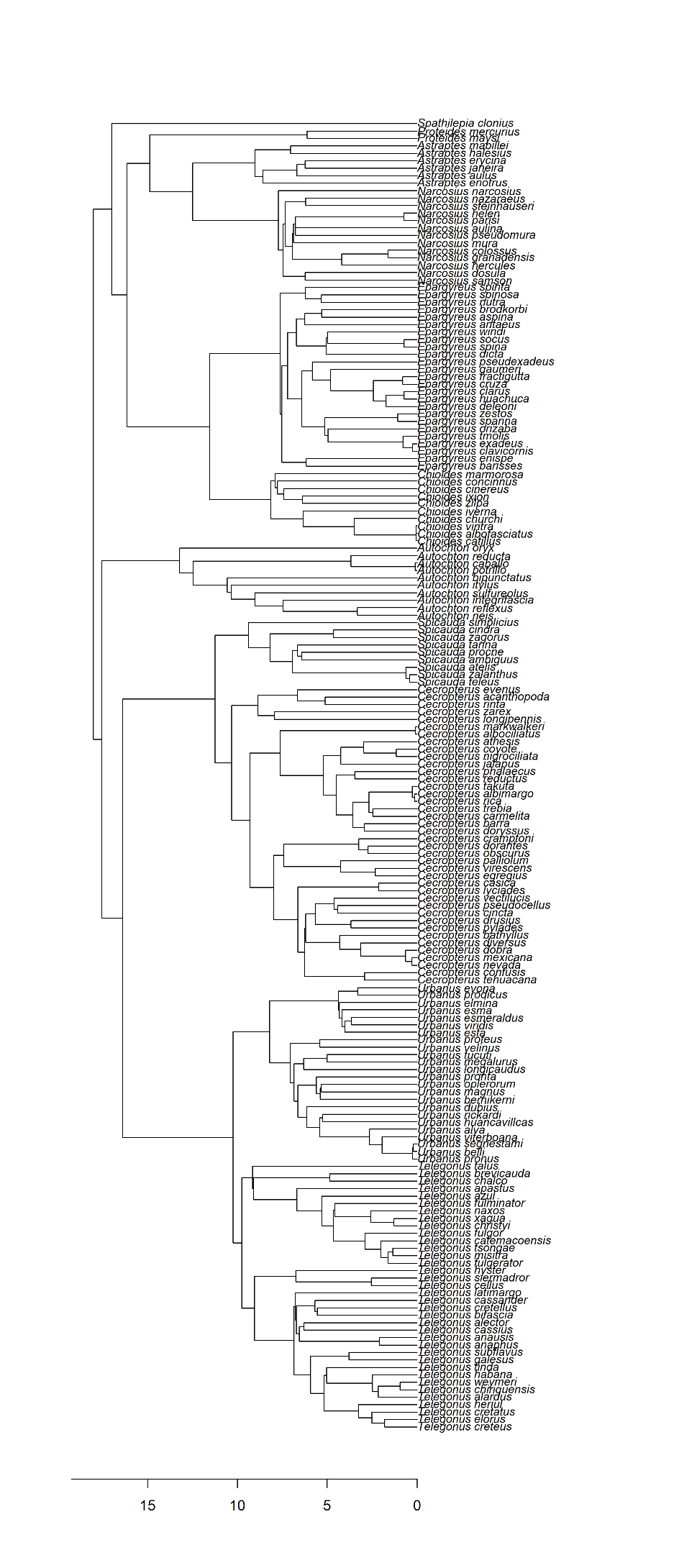

**Figure S1**: MCC tree used for phylogenetic analysis*.*

**Appendix S3:** Custom multivariate regression

```
function(Y,x,degree,plot.data=FALSE) {
	Y <- as.matrix(Y) ; x <- as.matrix(x)
	X <- 1 ; XX <- 1
	xx <- seq(min(x)-.05*(max(x)-min(x)),max(x)+.05*(max(x)-min(x)),length.out=200)
	for (i in 1:degree) { X <- cbind(X,x^i) ; XX <- cbind(XX,xx^i) }

	P <- solve(t(X)%*%X)%*%t(X)%*%Y
	predicted <- X%*%P
	residuals <- Y-predicted

	if ( isTRUE(plot.data) ) {
		plot(cbind(x,Y),pch=20,bty='l',xlab="Independent variable",ylab="Dependent variable 1")
		lines(cbind(xx,XX%*%P),col=2) ; #points(x,predicted[,1],pch=20,col=2,cex=.5)
		segments(x,predicted[,1],x,as.matrix(Y)[,1],col=2,lwd=.1)
	}

	F.ratio <- (sum(scale(predicted,scale=F)^2)/degree)/(sum(residuals^2)/(length(x)-degree-1))
	p.value <- pf(F.ratio, ncol(Y)*degree, ncol(Y)*(length(x)-degree-1),lower.tail=FALSE)
	R2 <- sum(scale(predicted,scale=F)^2)/sum(scale(Y,scale=F)^2)
	adjustR2 <- 1 - (1-R2)*((length(x)-1)/(length(x)-length(P)))

	p.R2 <- c()
	for (i in 1:9999) {
		per <- sample(1:length(x),length(x),replace=F)
		p.P <- solve(t(X)%*%X)%*%t(X)%*%Y[per,]
		p.predicted <- X%*%p.P
		p.residuals <- Y[per,]-p.predicted
		p.R2 <- c(p.R2,sum(scale(p.predicted,scale=F)^2)/sum(scale(Y[per,],scale=F)^2))
	}
	p.R2 <- (1+length(which(p.R2>=R2)))/10000


	Tab <- cbind(R2,adjustR2,F.ratio,ncol(Y)*degree, ncol(Y)*(length(x)-degree-1),p.value,p.R2)
	colnames(Tab) <- c("R^2","Adj. R^2","F-ratio","df1","df2","P(param)","P(permut)")
	print(Tab)
	list(predicted=predicted,residuals=residuals,param=P)
}
```

**Appendix S4:** Ancestral state reconstruction

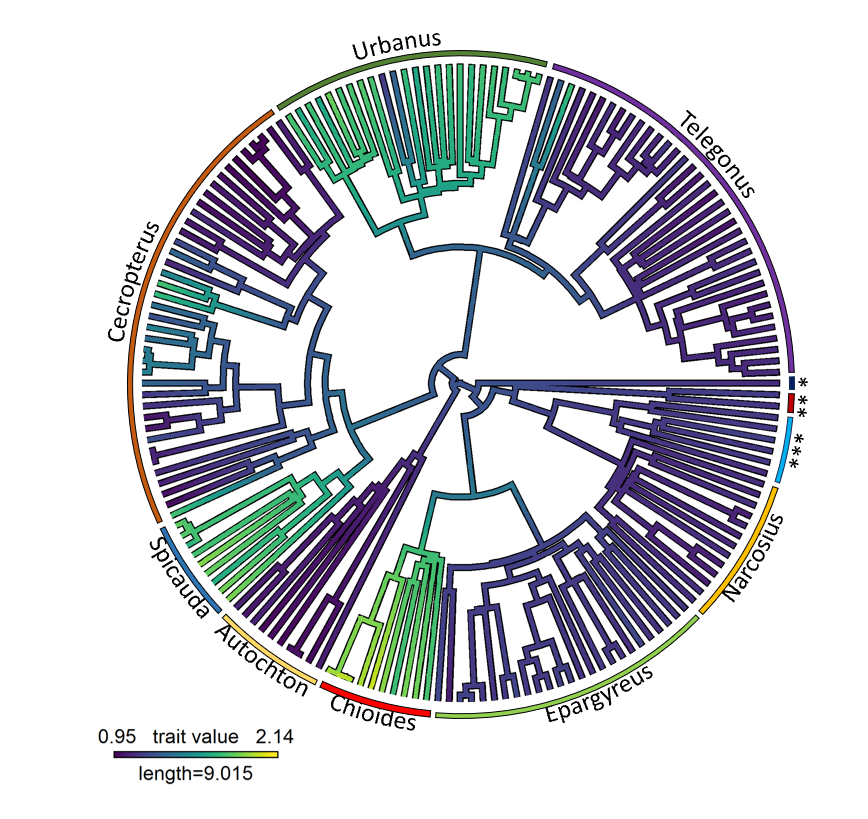

**Figure S3:** Ancestral state reconstruction for tail ratio across Eudamina. * Spathilepia, ** Proteides and *** Astraptes.

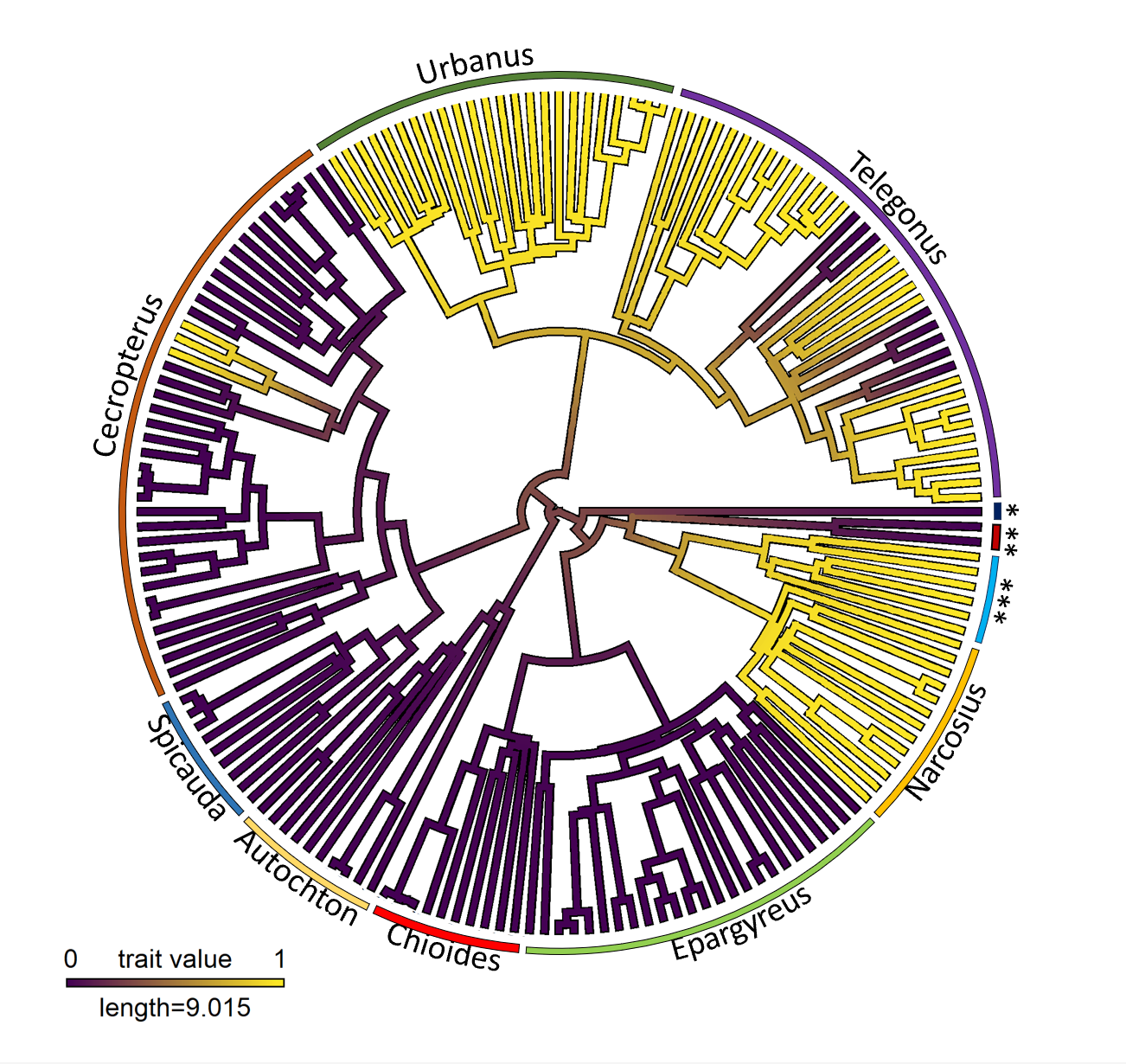

**Figure S4:** Ancestral state reconstruction for the presence or absence of dorsal iridescence across Eudamina. * Spathilepia, ** Proteides and *** Astraptes.

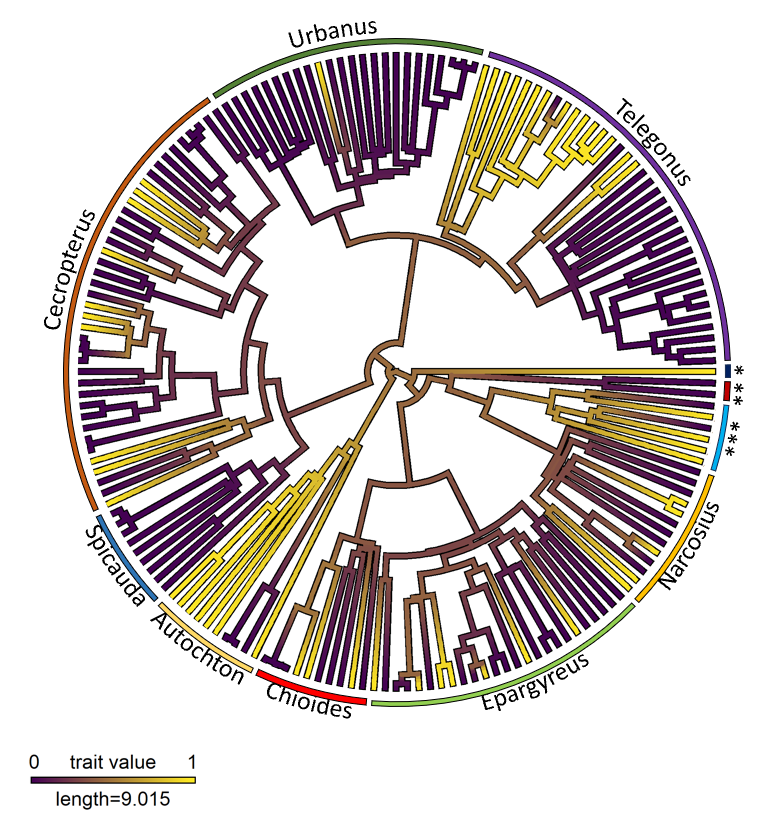
**Figure S5:** Ancestral state reconstruction for the presence or absence of creamy bands across Eudamina. * Spathilepia, ** Proteides and *** Astraptes.

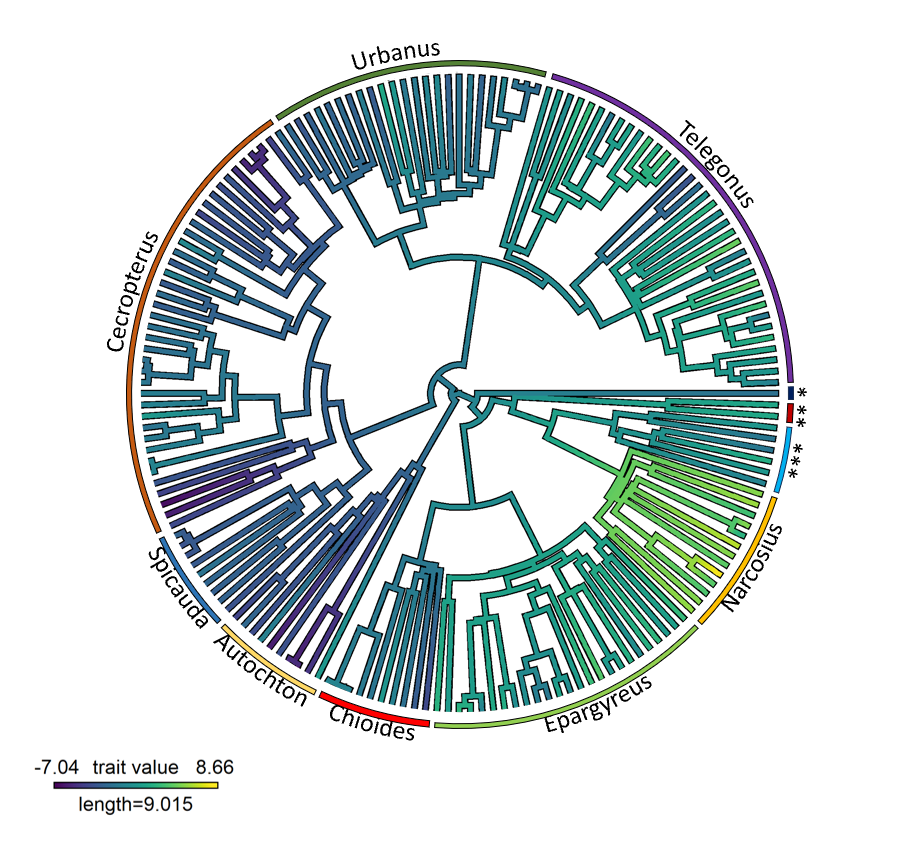
 **Figure S6:** Ancestral state reconstruction for size (PC1) across Eudamina. * Spathilepia, ** Proteides and *** Astraptes.

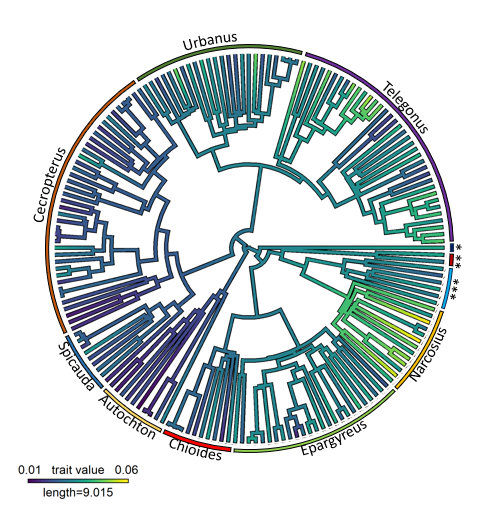

**Figure S7:** Ancestral state reconstruction for wing loading (based on fore wing area) across Eudamin. * Spathilepia, ** Proteides and *** Astraptes.

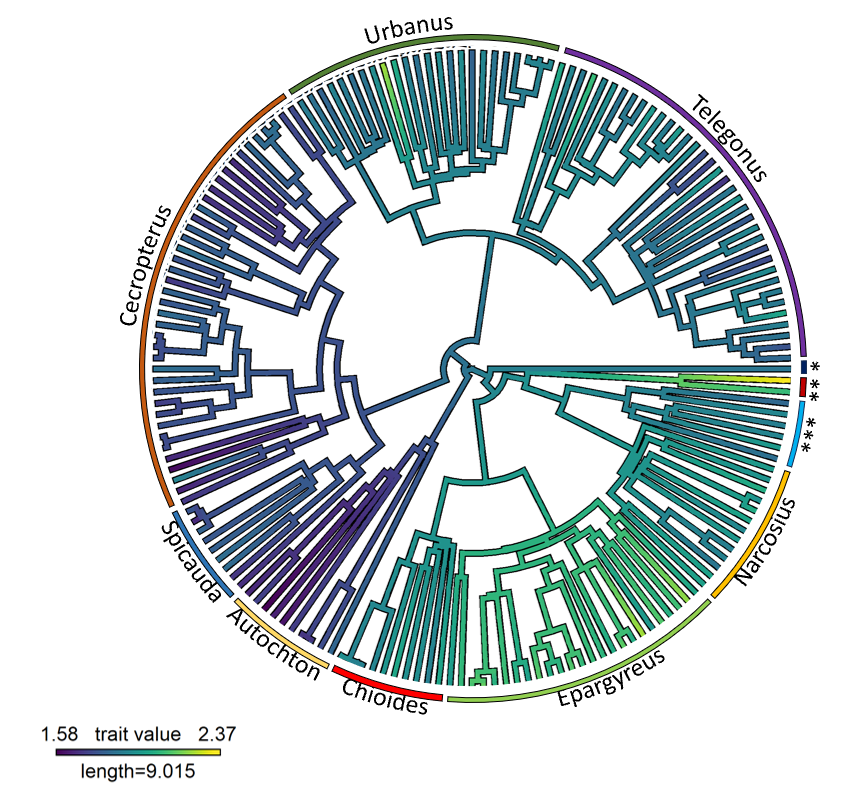

**Figure S8:** Ancestral state reconstruction for aspect ratio across Eudamina. * Spathilepia, ** Proteides and *** Astraptes.

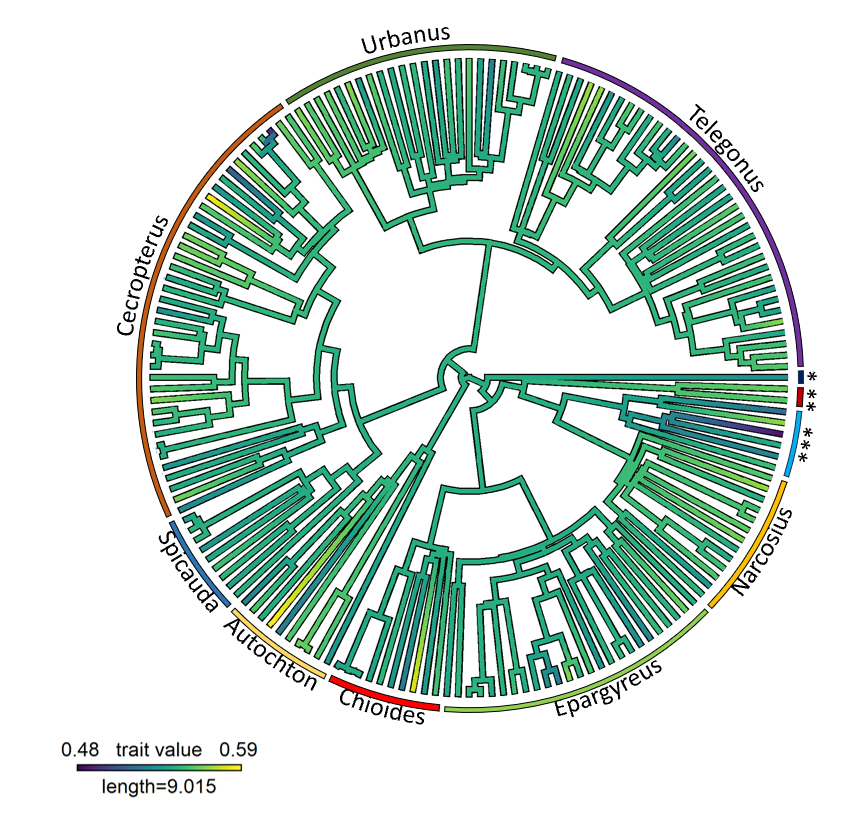
**Figure S9:** Ancestral state reconstruction for second moment of area across Eudamina. * Spathilepia, ** Proteides and *** Astraptes.

**Appendix S5:** Full brms outputs including diagnostic plots

*Model 1: Using size (PC1) as response*

```
Family: gaussian
  Links: mu = identity; sigma = identity
Formula: scale(PC1) ~ bands + iridescence + scale(tail_ratio) + distribution + (1 | gr(species_genetic, cov = GenDist))
   Data: factors (Number of observations: 176)
  Draws: 4 chains, each with iter = 4000; warmup = 1000; thin = 1;
         total post-warmup draws = 12000

Group-Level Effects:
~species_genetic (Number of levels: 176)
              Estimate Est.Error l-95% CI u-95% CI Rhat Bulk_ESS Tail_ESS
sd(Intercept)     0.18      0.02     0.15     0.21 1.00     1929     4288

Population-Level Effects:
                       Estimate Est.Error l-95% CI u-95% CI Rhat Bulk_ESS Tail_ESS
Intercept                 -0.48      0.29    -1.04     0.09 1.00     3854     6294
bandsyes                  -0.07      0.11    -0.28     0.16 1.00     7060     8716
iridescenceyes             0.30      0.22    -0.14     0.74 1.00     4994     7223
scaletail_ratio           -0.14      0.09    -0.31     0.04 1.00     4652     6825
distributionTropical       0.25      0.14    -0.02     0.52 1.00     6471     8510
distributionWidespread     0.44      0.22     0.01     0.88 1.00     6368     8704

Family Specific Parameters:
      Estimate Est.Error l-95% CI u-95% CI Rhat Bulk_ESS Tail_ESS
sigma     0.34      0.04     0.26     0.42 1.00     1443     2956

Draws were sampled using sample(hmc). For each parameter, Bulk_ESS
and Tail_ESS are effective sample size measures, and Rhat is the potential
scale reduction factor on split chains (at convergence, Rhat = 1).
```

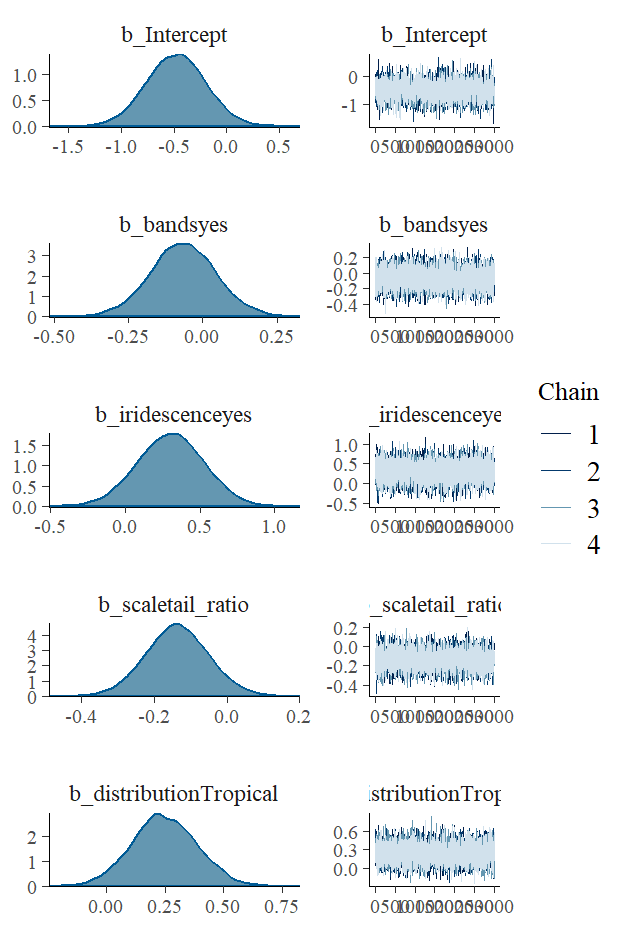

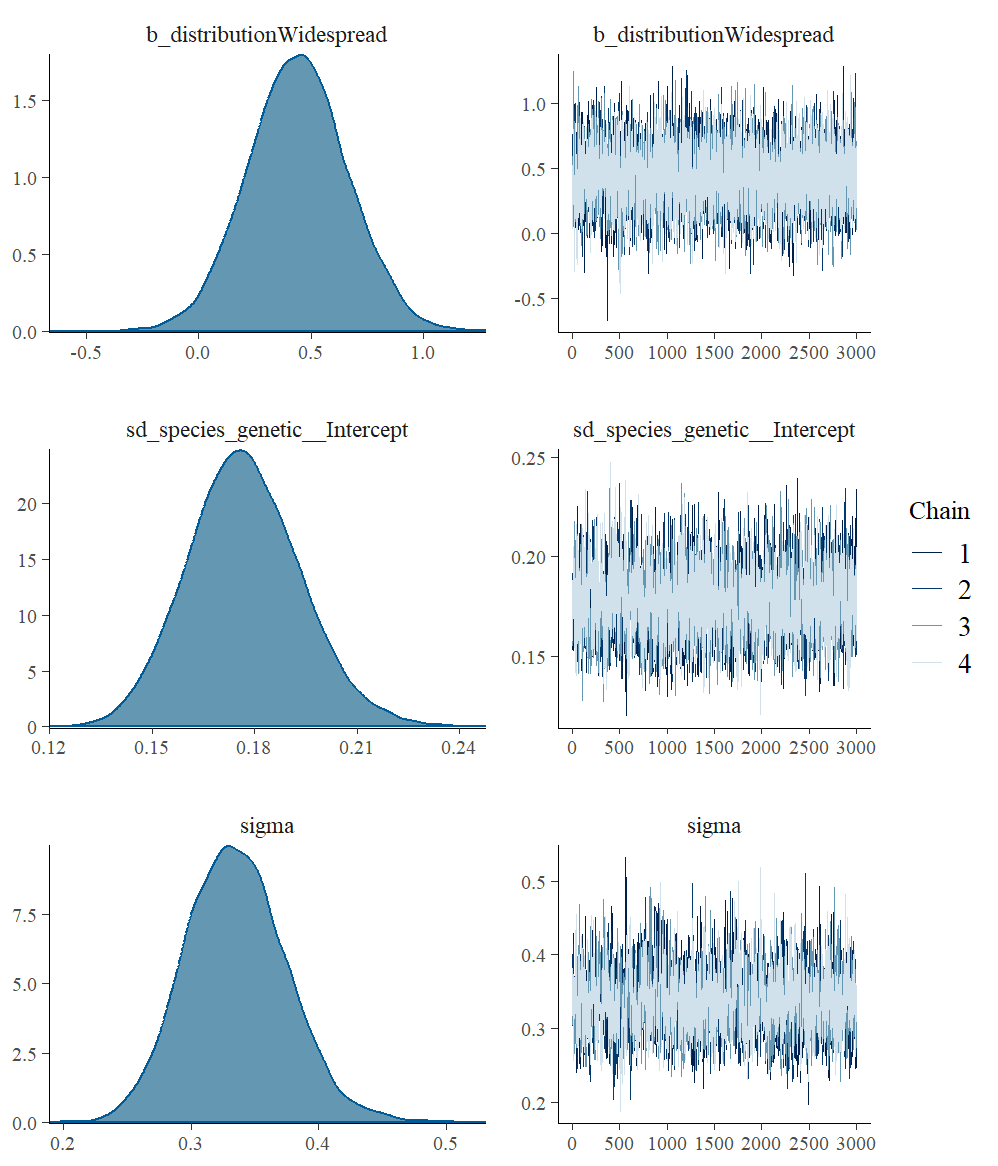

*Model 2: Using wing loading as a response.*

```
Family: gaussian
  Links: mu = identity; sigma = identity
Formula: scale(WL_F) ~ bands + iridescence + scale(tail_ratio) + distribution + (1 | gr(species_genetic, cov = GenDist))
   Data: factors (Number of observations: 176)
  Draws: 4 chains, each with iter = 4000; warmup = 1000; thin = 1;
         total post-warmup draws = 12000

Group-Level Effects:
~species_genetic (Number of levels: 176)
              Estimate Est.Error l-95% CI u-95% CI Rhat Bulk_ESS Tail_ESS
sd(Intercept)     0.15      0.03     0.10     0.21 1.00     2153     3718

Population-Level Effects:
                       Estimate Est.Error l-95% CI u-95% CI Rhat Bulk_ESS Tail_ESS
Intercept                 -0.48      0.30    -1.06     0.09 1.00     5999     7320
bandsyes                  -0.16      0.14    -0.44     0.12 1.00    10320     9487
iridescenceyes             0.94      0.24     0.46     1.41 1.00     7200     8342
scaletail_ratio           -0.19      0.09    -0.38    -0.01 1.00     6924     7818
distributionTropical       0.10      0.20    -0.30     0.49 1.00     9013     8392
distributionWidespread     0.41      0.30    -0.17     0.99 1.00     8534     9016

Family Specific Parameters:
      Estimate Est.Error l-95% CI u-95% CI Rhat Bulk_ESS Tail_ESS
sigma     0.64      0.06     0.53     0.75 1.00     2486     3996

Draws were sampled using sample(hmc). For each parameter, Bulk_ESS
and Tail_ESS are effective sample size measures, and Rhat is the potential
scale reduction factor on split chains (at convergence, Rhat = 1).
```

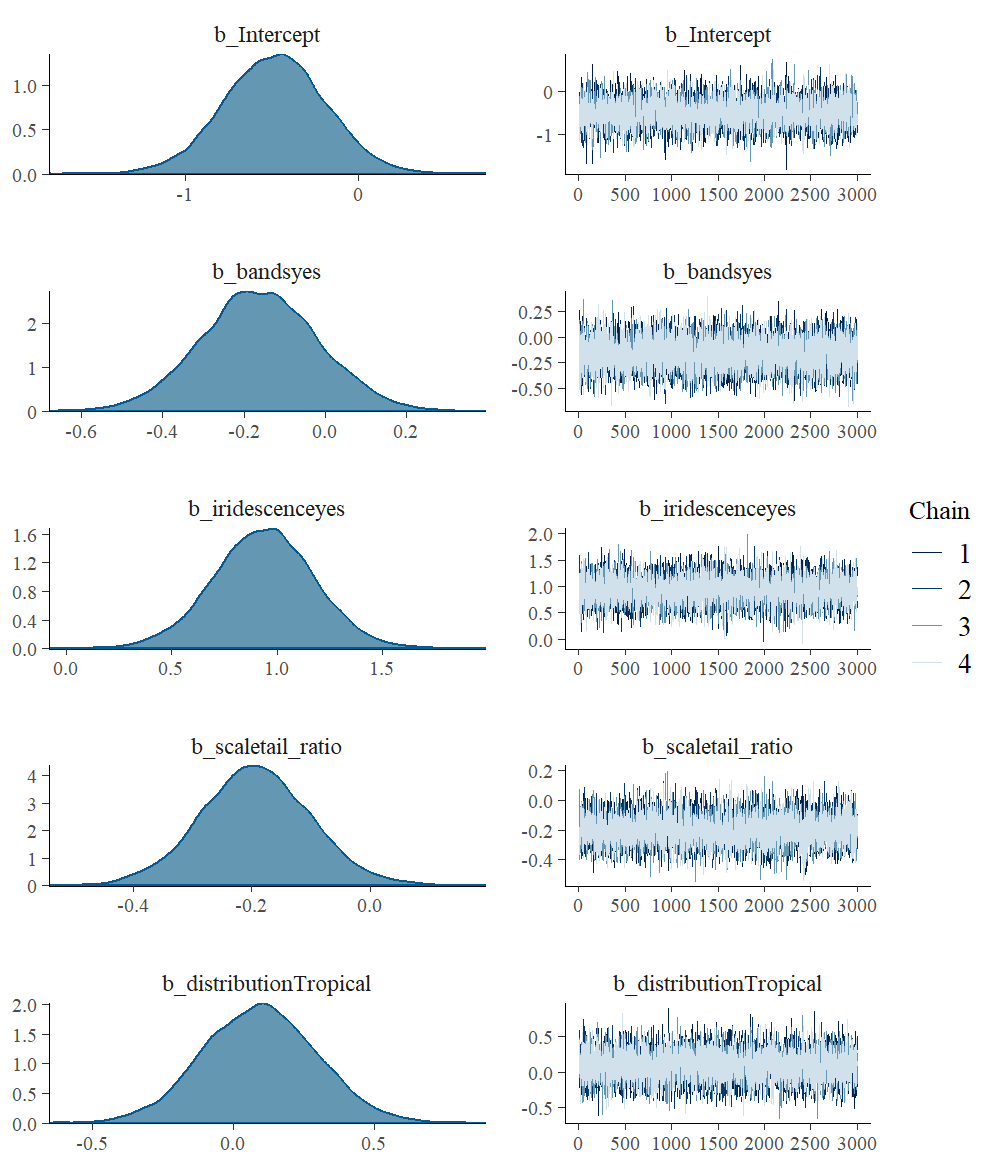

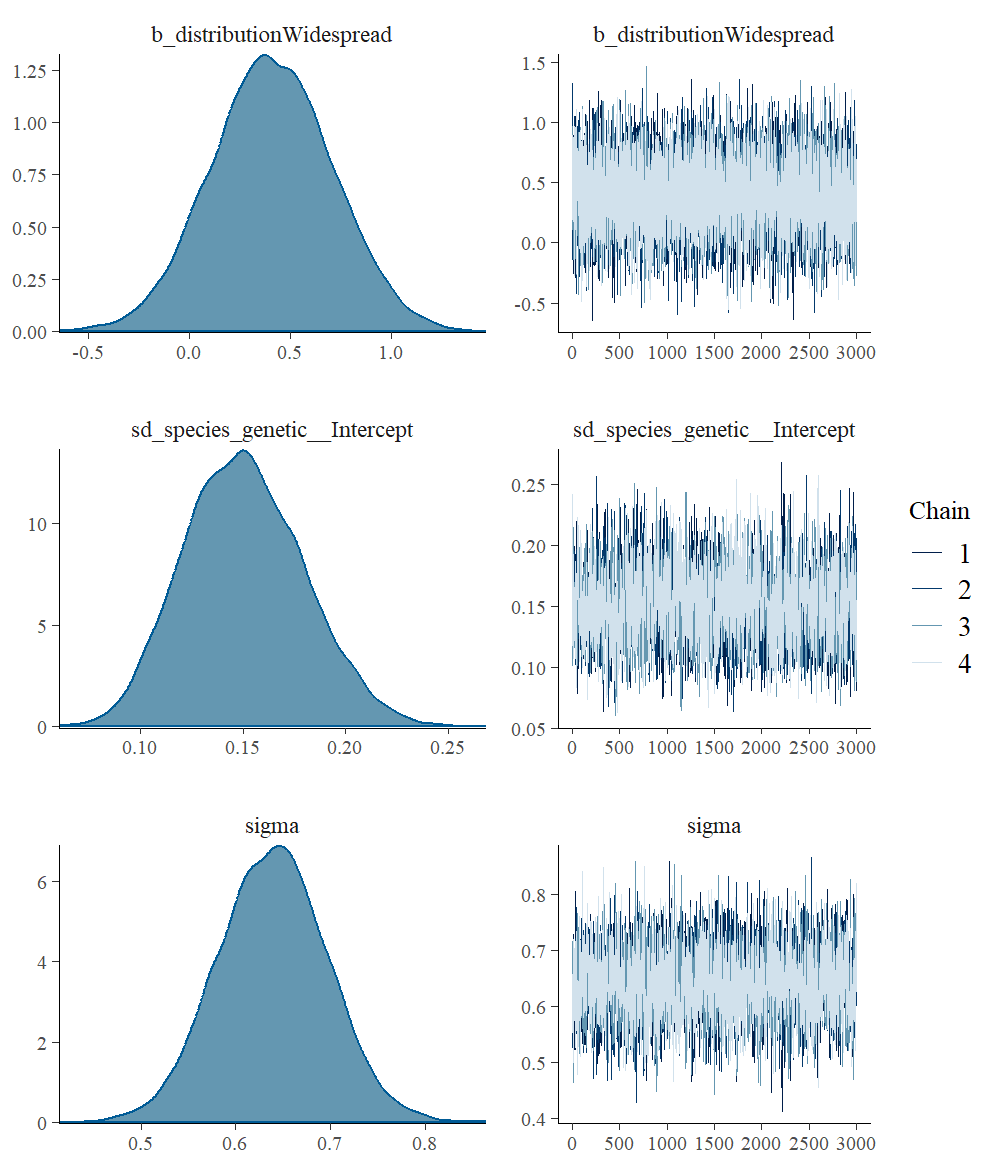
*Model 3: Using second moment of area as a response.*

```
Family: gaussian
  Links: mu = identity; sigma = identity
Formula: scale(non.dim.2nd_moment) ~ bands + iridescence + scale(tail_ratio) + distribution + (1 | gr(species_genetic, cov = GenDist))
   Data: factors (Number of observations: 176)
  Draws: 4 chains, each with iter = 4000; warmup = 1000; thin = 1;
         total post-warmup draws = 12000

Group-Level Effects:
~species_genetic (Number of levels: 176)
              Estimate Est.Error l-95% CI u-95% CI Rhat Bulk_ESS Tail_ESS
sd(Intercept)     0.05      0.03     0.00     0.13 1.00     2905     4816

Population-Level Effects:
                       Estimate Est.Error l-95% CI u-95% CI Rhat Bulk_ESS Tail_ESS
Intercept                 -0.33      0.24    -0.81     0.14 1.00    10665     9062
bandsyes                  -0.28      0.17    -0.61     0.06 1.00    16870     9525
iridescenceyes             0.12      0.19    -0.25     0.50 1.00    11241     7606
scaletail_ratio           -0.11      0.09    -0.29     0.06 1.00    13190     9604
distributionTropical       0.43      0.24    -0.05     0.90 1.00    11957     9100
distributionWidespread     0.26      0.36    -0.45     0.96 1.00    12816     8972

Family Specific Parameters:
      Estimate Est.Error l-95% CI u-95% CI Rhat Bulk_ESS Tail_ESS
sigma     0.99      0.06     0.88     1.10 1.00    10583     8057

Draws were sampled using sample(hmc). For each parameter, Bulk_ESS
and Tail_ESS are effective sample size measures, and Rhat is the potential
scale reduction factor on split chains (at convergence, Rhat = 1).
```

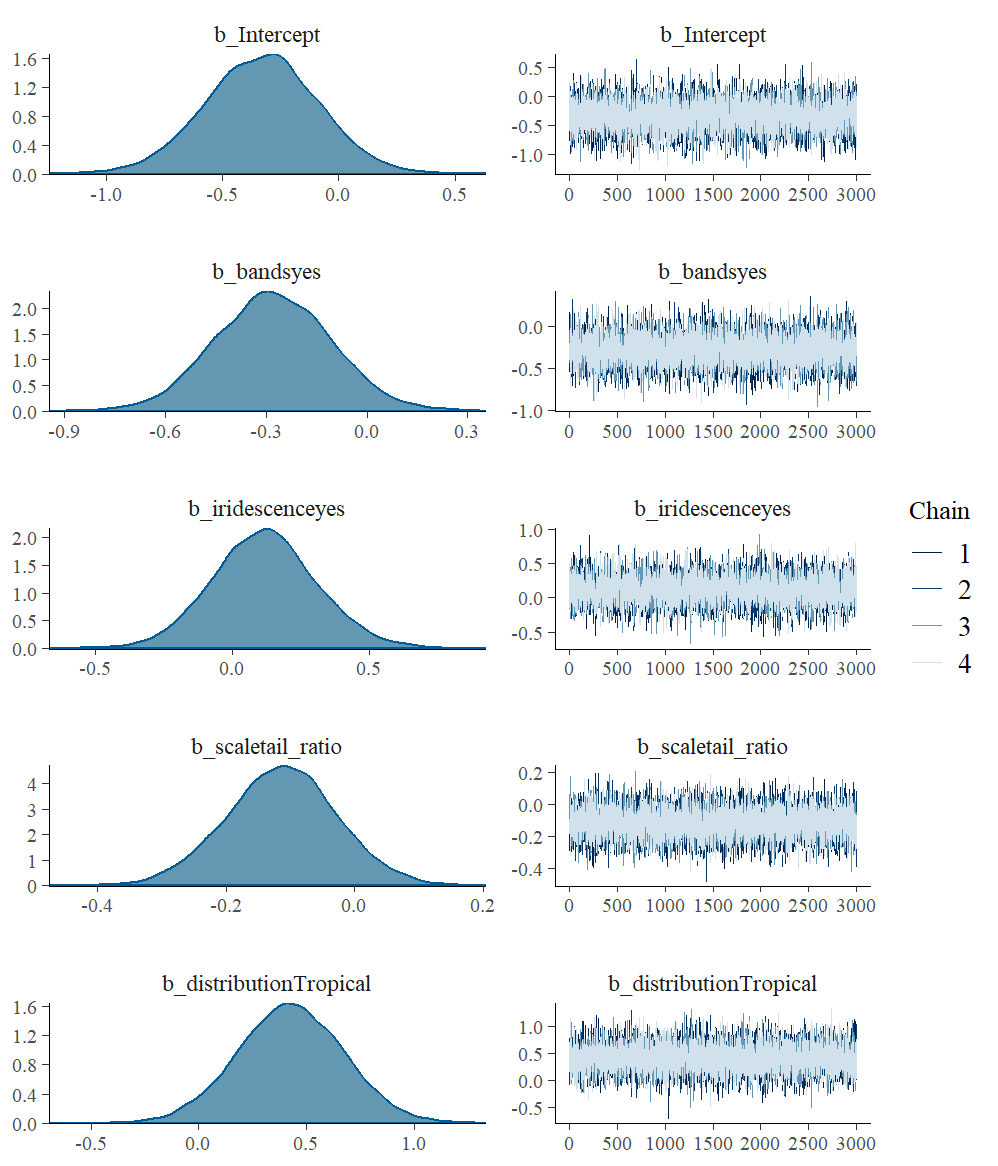

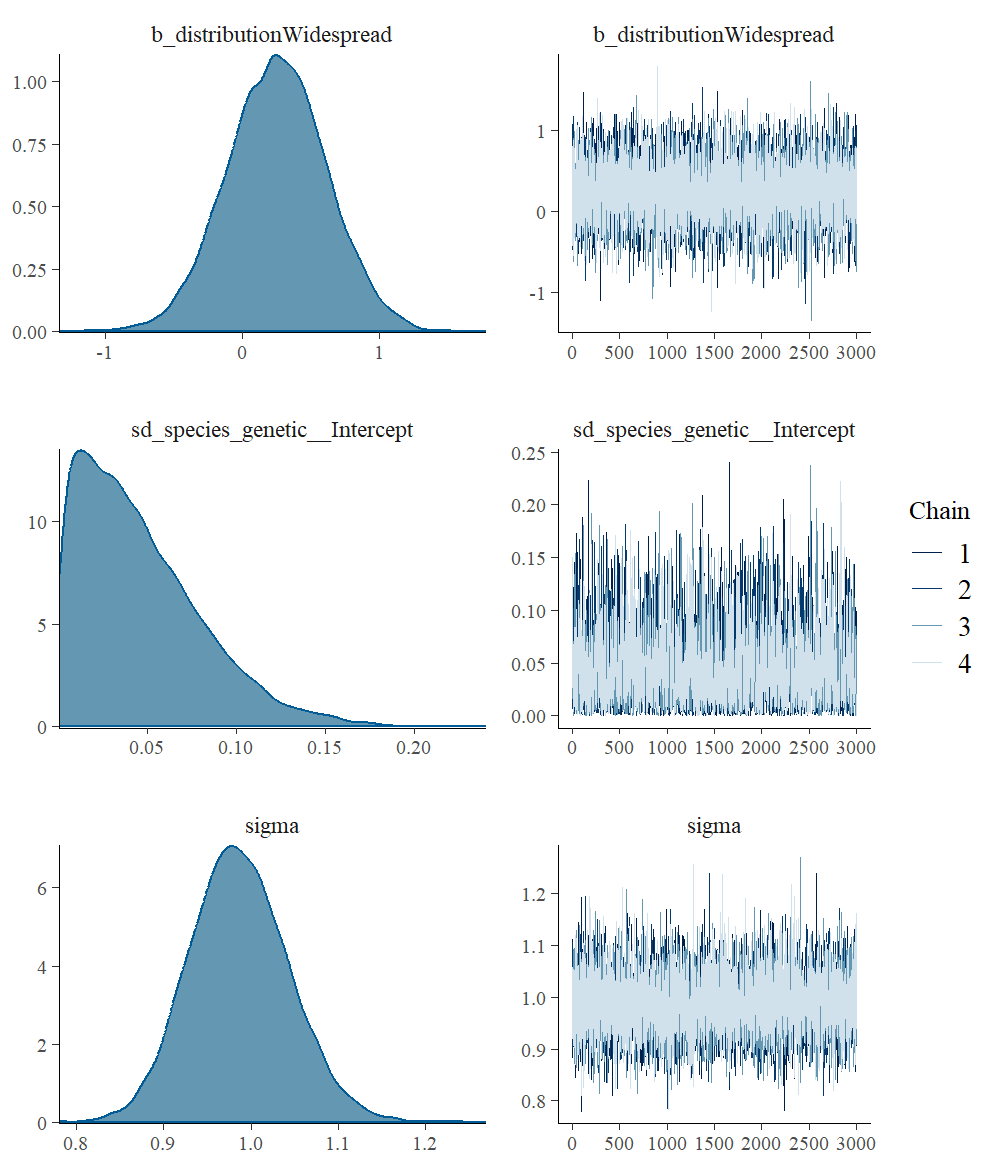

*Model 3: Using aspect ratio as a response.*

```
Family: gaussian
  Links: mu = identity; sigma = identity
Formula: scale(AR) ~ bands + iridescence + scale(tail_ratio) + distribution + (1 | gr(species_genetic, cov = GenDist))
   Data: factors (Number of observations: 176)
  Draws: 4 chains, each with iter = 4000; warmup = 1000; thin = 1;
         total post-warmup draws = 12000

Group-Level Effects:
~species_genetic (Number of levels: 176)
              Estimate Est.Error l-95% CI u-95% CI Rhat Bulk_ESS Tail_ESS
sd(Intercept)     0.18      0.02     0.15     0.21 1.00     2112     4380

Population-Level Effects:
                       Estimate Est.Error l-95% CI u-95% CI Rhat Bulk_ESS Tail_ESS
Intercept                 -0.28      0.29    -0.85     0.29 1.00     4494     5814
bandsyes                  -0.10      0.11    -0.32     0.13 1.00     6436     8428
iridescenceyes             0.49      0.23     0.05     0.93 1.00     6125     7044
scaletail_ratio           -0.01      0.09    -0.18     0.16 1.00     4785     6577
distributionTropical       0.07      0.14    -0.21     0.34 1.00    10462     9219
distributionWidespread     0.22      0.23    -0.22     0.66 1.00     7835     8715

Family Specific Parameters:
      Estimate Est.Error l-95% CI u-95% CI Rhat Bulk_ESS Tail_ESS
sigma     0.35      0.04     0.27     0.43 1.00     1680     2857

Draws were sampled using sample(hmc). For each parameter, Bulk_ESS
and Tail_ESS are effective sample size measures, and Rhat is the potential
scale reduction factor on split chains (at convergence, Rhat = 1).
```

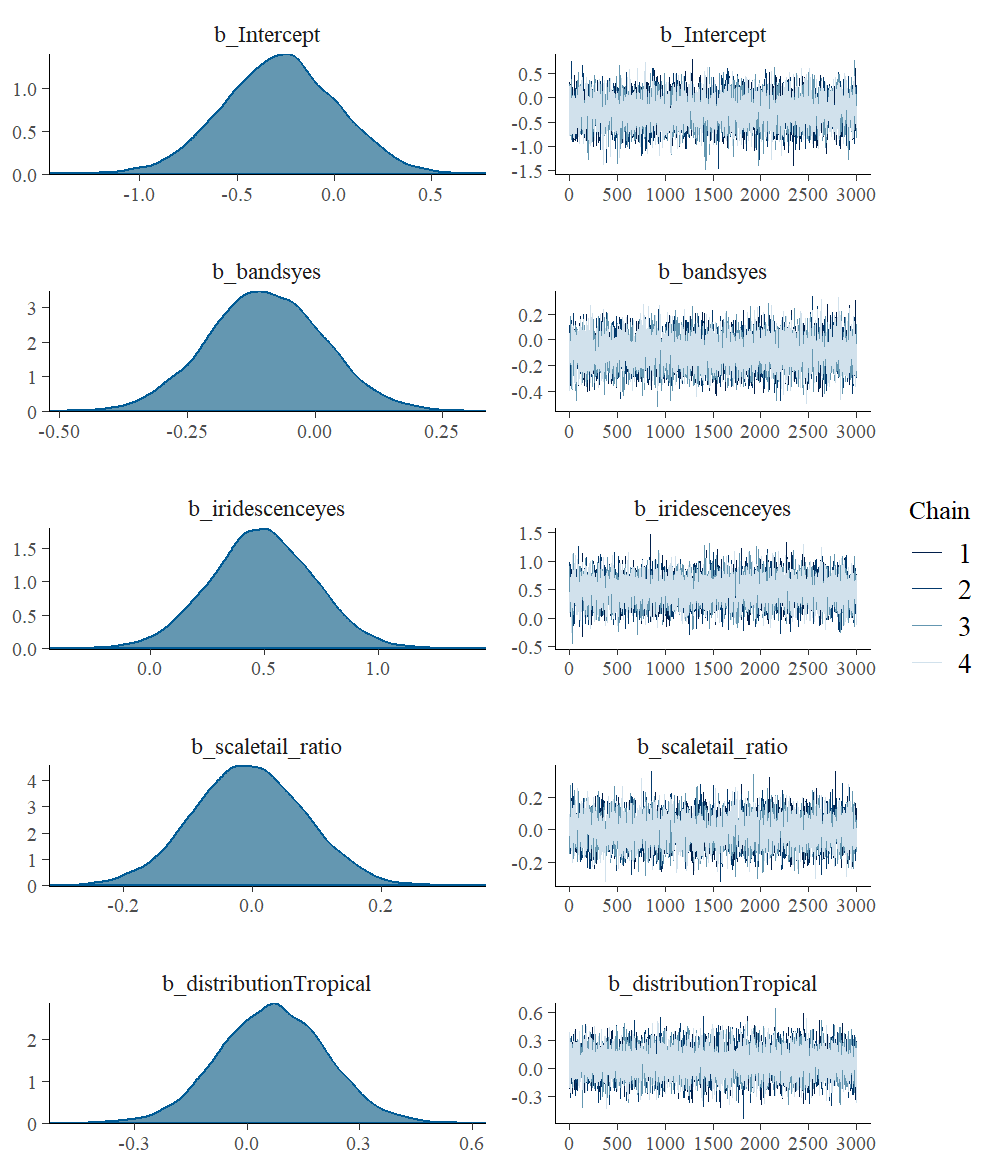

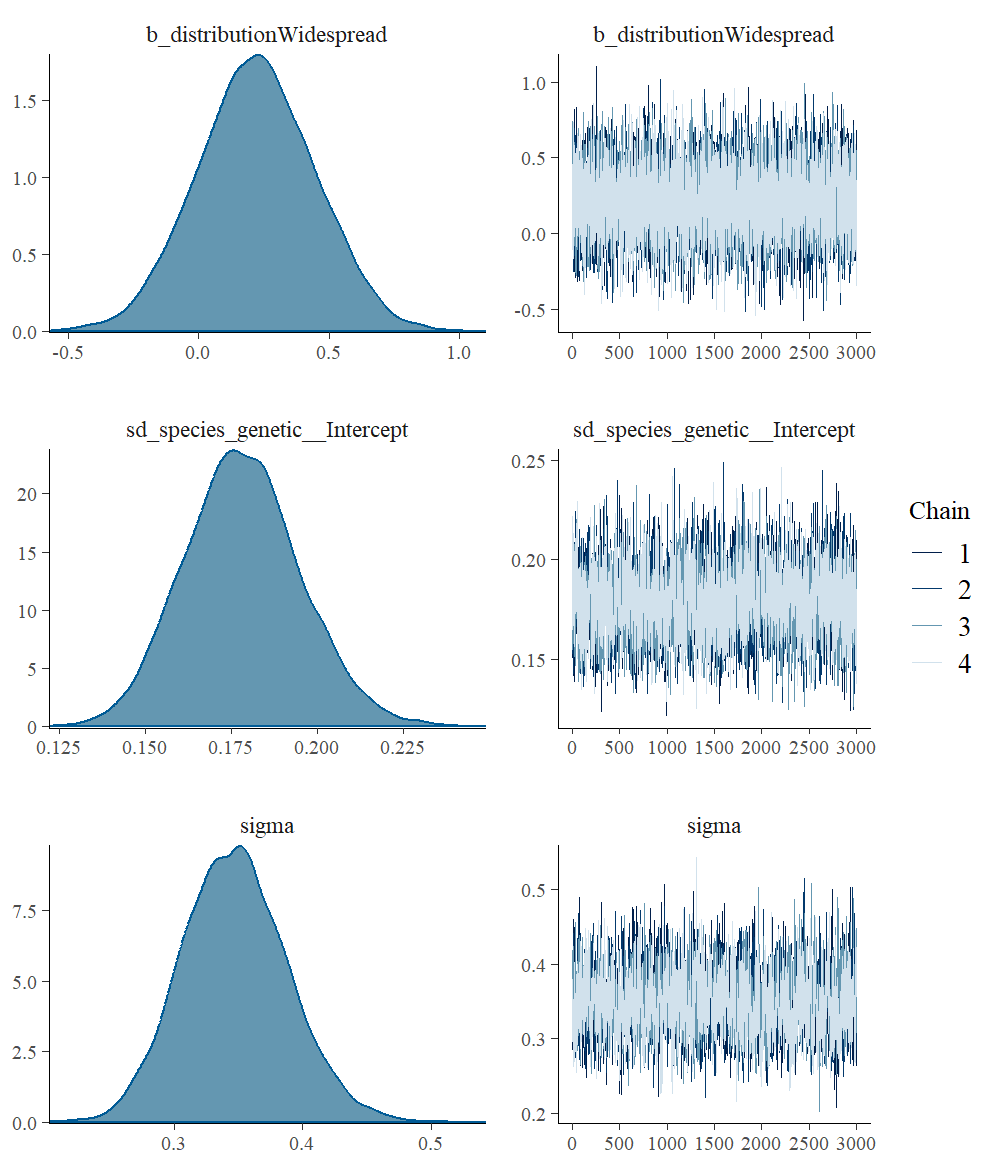

**Appendix S6:** Phylogenetic PLS analysis

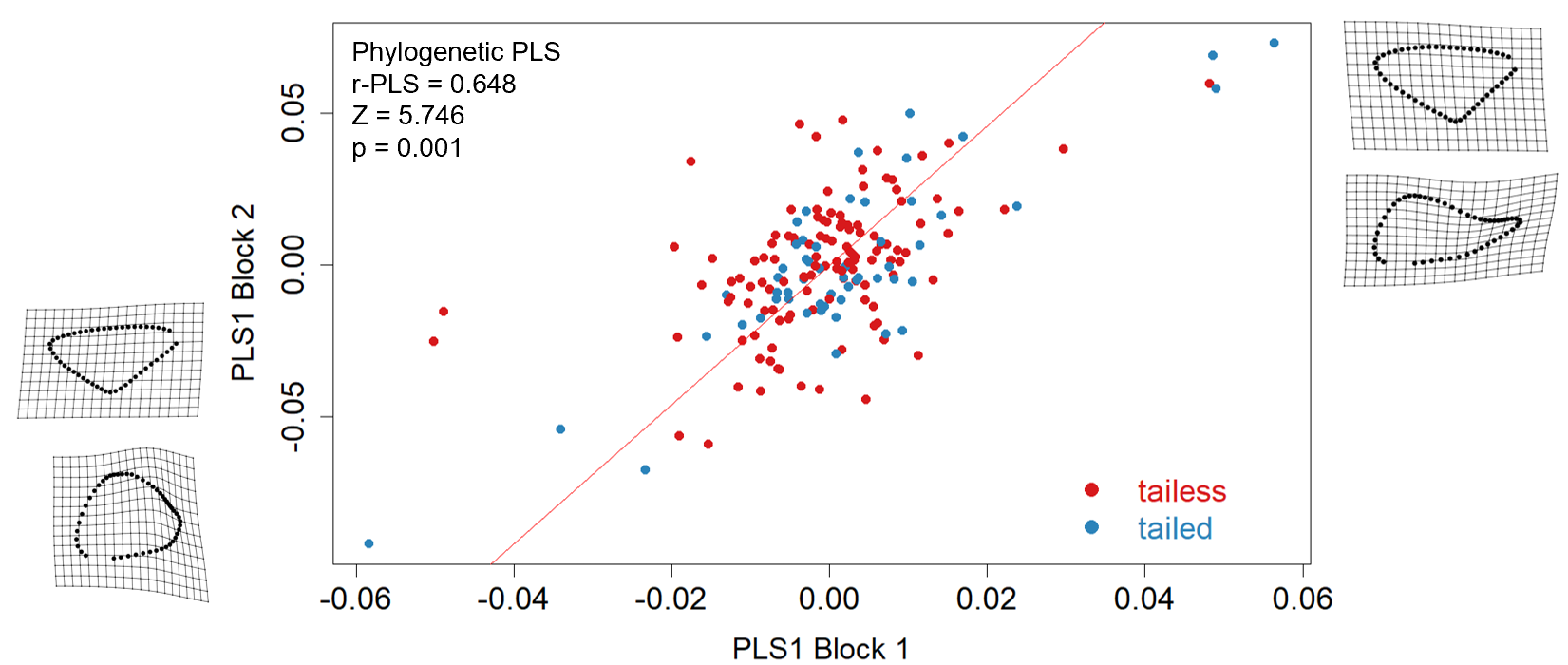

**Figure S10:** Phylogenetic PLS analysis showing covariation between forewing and hindwing shapes. Wing shapes (forewings on x-axis, hindwings on y-axis) in the corner represent expected shapes in the extremes of their variation. *Chioides vintra* was excluded from this analysis due to its unusually short genetic distance from its sister species, Chioides catillus, combined with an extremely large phenotypic divergence between both species. Colour indicates the presence and absence of hindwing tails (presence = tail ratio > 1.3).
